# Evolution of gene expression across species and specialized zooids in Siphonophora

**DOI:** 10.1101/2021.07.30.454354

**Authors:** Catriona Munro, Felipe Zapata, Mark Howison, Stefan Siebert, Casey W. Dunn

## Abstract

**Background:** Siphonophores are complex colonial animals, consisting of asexually-produced bodies (called zooids) that are functionally specialized for specific tasks, including feeding, swimming, and sexual reproduction. Though this extreme functional specialization has captivated biologists for generations, its genomic underpinnings remain unknown. We use RNA-seq to investigate gene expression patterns in five zooids and one specialized tissue (pneumatophore) across seven siphonophore species. Analyses of gene expression across species present several challenges, including identification of comparable expression changes on gene trees with complex histories of speciation, duplication, and loss. Here, we conduct three analyses of expression. First, we examine gene expression within species. Then, we conduct classical analyses examining expression patterns between species. Lastly, we introduce Speciation Branch Filtering, which allows us to examine the evolution of expression in a phylogenetic framework.

**Results:** Within and across species, we identified hundreds of zooid-specific and species-specific genes, as well as a number of putative transcription factors showing differential expression in particular zooids and developmental stages. We found that gene expression patterns tended to be largely consistent in zooids with the same function across species, but also some large lineage-specific shifts in gene expression.

**Conclusions:** Our findings show that patterns of gene expression have the potential to define zooids in colonial organisms. We also show that traditional analyses of the evolution of gene expression focus on the tips of gene phylogenies, identifying large-scale expression patterns that are zooid or species variable. The new explicit phylogenetic approach we propose here focuses on branches (not tips) offering a deeper evolutionary perspective into specific changes in gene expression within zooids along all branches of the gene (and species) trees.

## Background

Colonial animals, consisting of genetically identical bodies that are physically and physiologically connected, can be found across the metazoan tree of life [1]. Functional specialization of such bodies has evolved multiple times within colonial animals, with siphonophores in particular showing the greatest diversity of functionally specialized bodies [2]. Siphonophores are highly complex, colonial “superorganisms” consisting of asexually produced bodies (termed zooids) that are homologous to solitary free-living polyps and medusae (the typical body forms in Cnidaria), but that share a common gastrovascular cavity [3–7]. The extreme specialization of siphonophore zooids has been of central interest to zoologists since the 19th century, in part because these zooids are highly interdependent [2,4]. Siphonophore zooids have been likened to animal organs, with each functionally specialized zooid performing distinct roles within the colony [4]. While this analogy works well for understanding the function of zooids within the colony as a whole, the analogy falls short in terms of explaining the evolutionary origin and development of these biological units: functionally specialized zooids are evolutionarily homologous to free living organisms, they are multicellular, possess distinct zooid specific cell types, and show regional subfunctionalization [7–9]. While the developmental mechanisms generating zooids are very different in different clades of colonial animals [3,10–13], the evolutionary processes acting on zooids may be similar to those acting on other modular biological units such as cell type, tissue, and organ [1]. The cellular and molecular processes underlying the patterning and molecular function of functionally specialized cnidarian zooids remain an open biological question. In recent years there has been a focus in particular on differential gene expression patterns found in different functionally specialized zooids [14–16].

Efforts to investigate the functional specialization of siphonophores have been limited, in part because there have been few detailed investigations of zooid structure. In the last half century, the microanatomy of siphonophore zooids and tissues has been investigated in only a handful of siphonophore species [8,9,11,17]. This leaves many unknowns about how zooid structure and function differ across zooid types and species. Recent *in situ* gene expression analyses in siphonophores have described where a small number of preselected genes are expressed at high spatial resolution [8,13,16], but these methods are limited since they require a large number of specimens per gene and siphonophores are relatively difficult to collect. RNA-seq analyses of hand-dissected specimens [14–16,18], in contrast, can describe the expression of a very large number of genes at low spatial resolution. The fact that so much data is obtained from each specimen is particularly advantageous in difficult-to-collect organisms like siphonophores. An earlier RNA-seq study of gene expression in two zooid types in a single siphonophore species showed the potential of this method to better understand differences between zooids [16].

In this study we use RNA-seq data to investigate the functional specialization and evolution of zooids across siphonophore species. We address this question at three comparative levels, namely within species, between species, and across species incorporating an explicit phylogenetic approach. To carry out these analyses, we also address three key challenges to study the evolution of gene expression in a comparative framework [19]. First, we introduce a new metric to normalize gene expression which is valid for comparison across species. A common metric used to quantify gene expression is Transcripts Per Million (TPM) [20,21]. TPM is a relative measure of expression which depends on the number of genes present in a reference. Hence, TPM values are a valid metric when comparing libraries within a single species, but are not directly comparable across species when the references are incomplete, or genes have been gained and lost in the course of evolution. To address this issue, we introduce Transcripts Per Million 10K (TPM10K), a metric which normalizes TPM to account for different sequencing depths among species (see Methods for details). We use TPM10K in our between species and phylogenetic-based analyses.

Second, we further account for gene and species effects to compare gene expression across genes and species. Normalized read counts produced by RNA-seq experiments are proportional to gene expression, but they are also impacted by factors that can vary across genes and species such as gene sequence and length. Therefore, to estimate and compare expected gene counts it is necessary to model such factors with unknown species- and gene-specific counting-efficiency coefficients [19]. A direct comparison of expected gene counts among species, however, can be misleading if differences in counts are simply due to differences in counting efficiency and not due to differences in expression. To address this issue, we use ratios of expected counts. Using ratios of counts, we are able to eliminate species- and gene-specific counting efficiency within species, as the unknown counting efficiency factor is in both the numerator and the denominator, and is thus removed prior to comparisons among species [19]. We use ratios of expected counts in our between species and phylogenetic-based analyses.

Third, we use a novel phylogenetic approach to investigate the evolution of gene expression on gene trees with complex evolutionary histories. Most evolutionary studies of gene expression focus exclusively on comparing expression values of strict orthologs (i.e., gene lineages related only by speciation events) which are shared across all species. Analyses of strict orthologs are limited to a subset of genes that have no evidence of duplication, and we call this approach Speciation Tree Filtering (STF) [22–25]. This is an all or nothing approach, where gene trees or subtrees (clades) are retained if they have one sequence for each species, and discarded if there are multiple sequences in any of the species. Because this is usually such a small fraction of genes, traditional STF analyses discard a large fraction of the data, including many gene families that may be of interest to the investigator.

Because the history of genes is characterized by more complex scenarios involving gene duplication and speciation, we introduce a new method that takes advantage of this rich history to examine the evolution of gene expression across more genes and species. We call this method Speciation Branch Filtering (SBF) (Fig. 2), as we map expression data to gene phylogenies and identify equivalent branches in the species tree which are descended from speciation events in order to make valid comparisons across species. While STF is a gene tree filtering approach, where entire trees or subtrees are discarded due to even a single duplication event, SBF is a branch filtering approach. This means that SBF retains many informative branches from trees that would be removed by STF, and preserves many more evolutionary comparisons for analyses. We use SBF in our explicit phylogenetic analyses across species.

**Figure 1:**
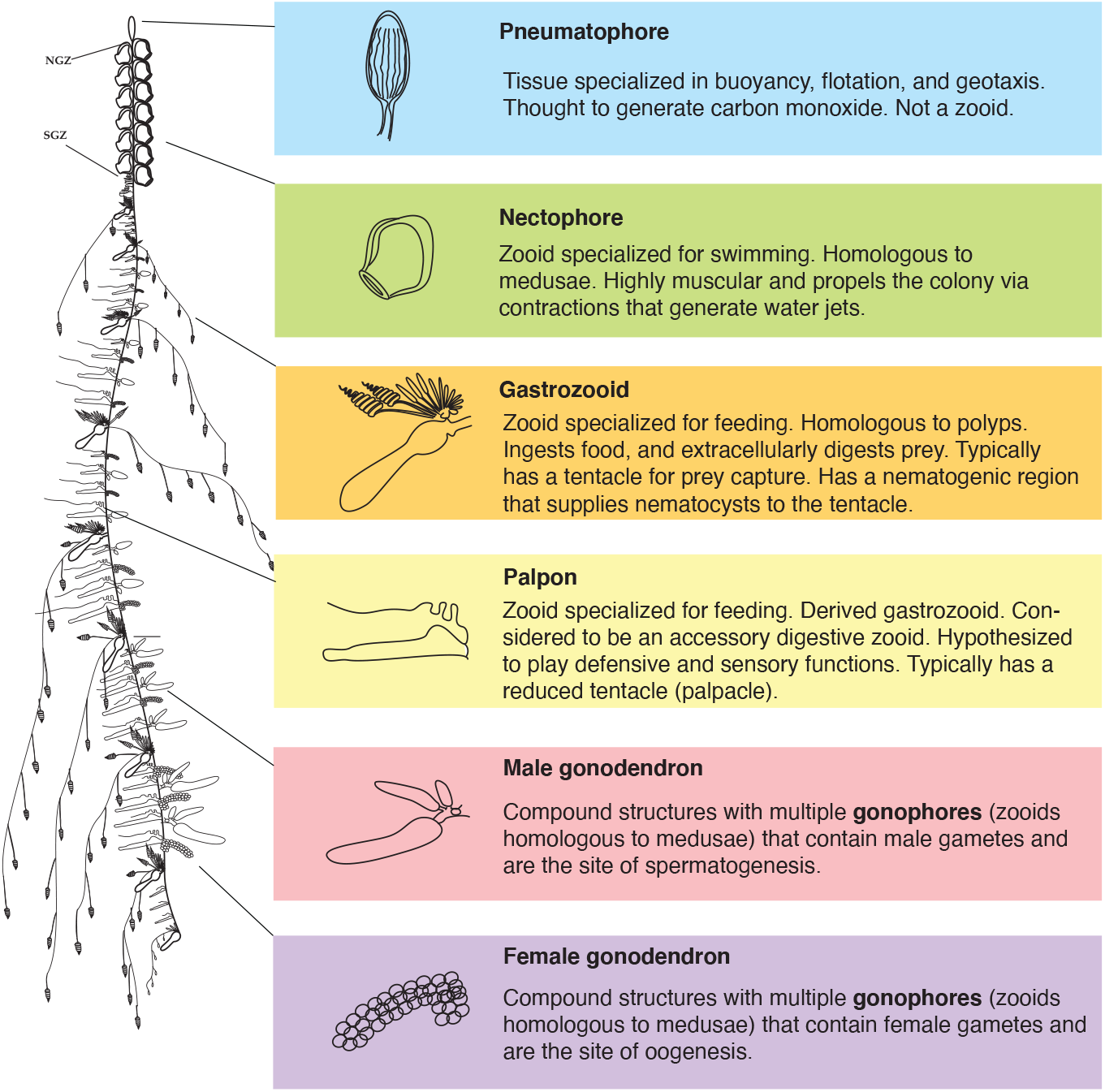
Schematic of a siphonophore colony, zooid organization, and zooid function. Here, we illustrate the siphonophore *Nanomia bijuga*, with highlighted zooids and pneumatophore, and explanations of known function. NGZ - nectosomal growth zone. SGZ - siphosomal growth zone. Diagram by Freya Goetz (http://commons.wikimedia.org/wiki/File:Nanomia_bijuga_whole_animal_and_growth_zones.svg)

**Figure 2:**
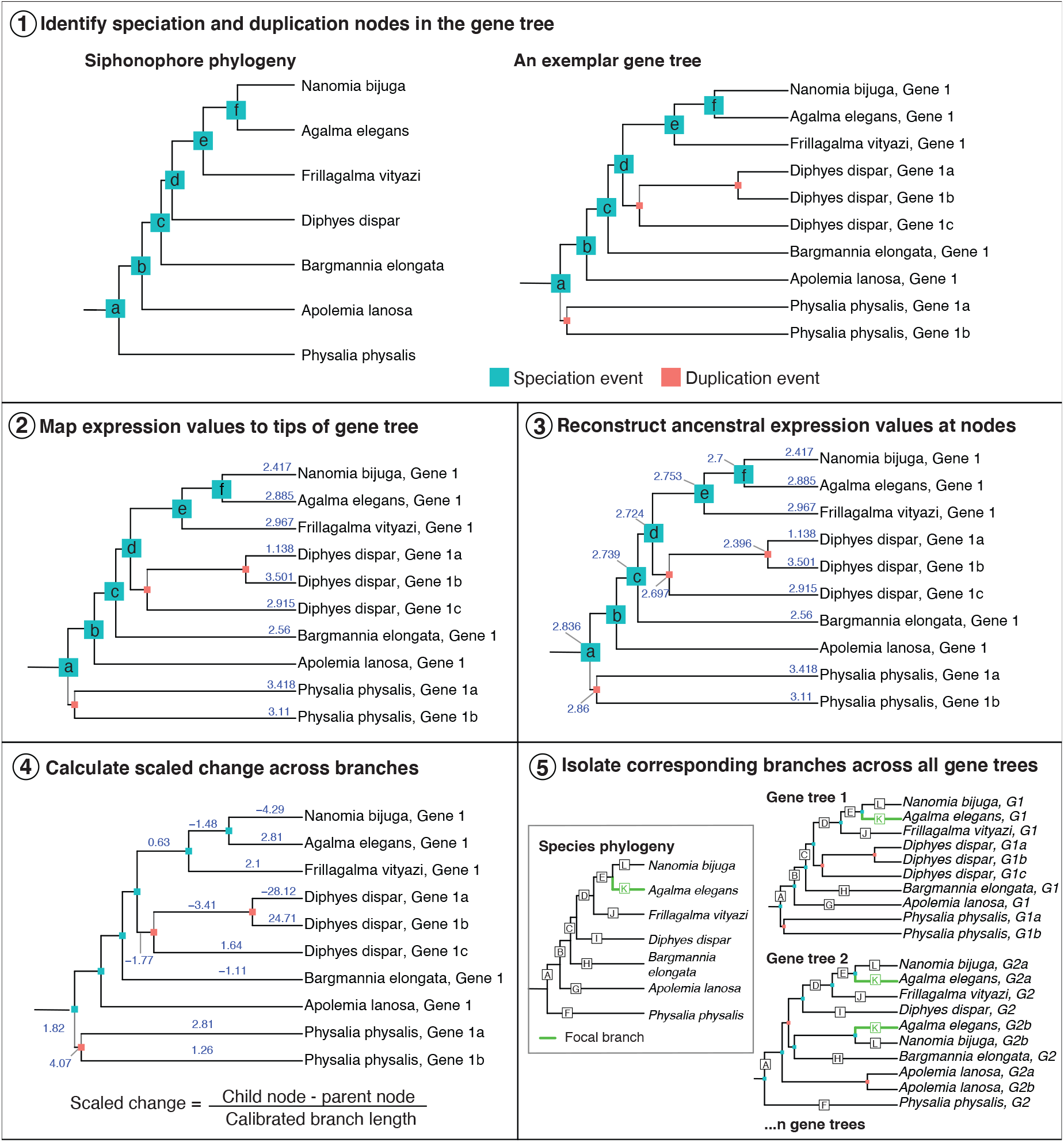
New phylogenetic approach to identify evolutionary changes in gene expression, called Speciation Branch Filtering (SBF). Step 1, we label each of the nodes in the species tree, and identify equivalent speciation nodes across every gene tree (an exemplar is shown here). Step 2, we map expression values (TPM10K) to the tips (expression values are mapped and reconstructed for each homologous zooid separately). Step 3, we reconstruct ancestral trait expression values at all internal nodes where expression data are available. Step 4, we calculate scaled change in gene expression (child node expression - parent node expression / branch length). Branch length is calibrated to the species tree branch lengths. Step 5, we identify branches in gene trees that correspond to equivalent branches in the species tree. There may be more than one branch in a gene tree that corresponds to the same branch in the species tree.

Briefly, the principles of SBF are as follows: first we identify speciation and duplication events in a given gene tree, specifically labelling nodes that correspond to speciation events in the species tree (Fig. 2, step 1). Next, we map gene expression values to the tips of the gene tree (Fig. 2, step 2), and use phylogenetic methods to reconstruct expression values at the internal nodes (Fig. 2, step 3). Expression values are mapped and reconstructed for each zooid/tissue separately, with the assumption that the structure is homologous across species. Then, we calculate scaled expression values across branches, by subtracting the expression value at the child node from the parent node, and dividing by branch length (branch lengths are calibrated against branches in the species tree) (Fig. 2, step 4). Finally, we identify branches in the gene trees that correspond to equivalent branches in the species tree (hereafter species-equivalent branches) (Fig. 2, step 5). Species-equivalent branches are gene tree branches with parent and child nodes that are both speciation events and that correspond to branches in the species tree. Each branch in the species tree is given a unique identifier (i.e., a letter) and the species-equivalent branches in gene trees are given the same identifier (Fig. 2, step 5). This method enables the selection of branches within gene trees that are equivalent to branches within the species tree, and are thus comparable with one another across all gene trees. Unlike the STF approach, this approach considers equivalent branches that are descended from speciation events, but that have more complex evolutionary histories. For example, due to deeper gene duplication events, gene trees often contain multiple branches that correspond to the same branch in the species tree (Fig. 2, step 5). Our method allows us to consider all of these branches.

Using the methods described above, combined with classical differential gene expression (DGE) analysis, we conducted DGE analyses among siphonophore zooids, and one specialized tissue (the pneumatophore, a gas filled float), in seven siphonophore species to address three questions. First, we analyzed differences in gene expression within species to identify patterns that define particular zooid types, and gain insight into the genes that may be playing a role in the structure, maintenance, and functioning of zooids. We also examined whether gene expression patterns could distinguish novel, distinct species-specific zooid types within particular species. Second, we compared gene expression between species, using STF followed by linear models to identify zooid- and species-variable genes. And third, we used phylogenetic methods and SBF to identify putative clade/lineage or zooid specific expression patterns.

## Results

### Within species analyses: which genes are specific to particular zooid types?

We sequenced mRNA from microdissected zooids from seven different siphonophore species (at least two replicate colonies) and mapped these short-read libraries to previously published transcriptomes [26]. We collected RNA-seq data from, where possible, 5 different zooids and one specialized tissue, the pneumatophore, as well as unique zooids specific to *Agalma elegans* (B palpons), *Physalia physalis* (tentacular palpon), and *Bargmannia elongata* (yellow and white gastrozooids), and where possible developing and mature zooids (Table S1). Due to species availability, we were not able to sample more than one replicate for some zooids - single replicates were excluded from downstream DGE analyses (see Fig. S1).

The first component of variation that we assessed was among technical replicates. The technical replicates consist of re-sequenced developing nectophores and developing gastrozooids from the same *Frillagalma vityazi* individual that were spiked in across multiple lanes and runs. Lane and run effects have been proposed as major sources of technical variability in RNA-seq data that may confound observations of biological variation [27,28]. The differences between technical replicates (Fig. S2) were much smaller (0.39% variance of expression distance) than the differences between zooids (98.32% variance of expression distance). Differences among technical replicates of the same zooid were correlated with library size and run, not by lane.

The second component of variation we considered was biological variation among sampled colonies (Fig. S3–S9). Specimens were collected in the wild at different depths and over different time periods, but despite these environmental factors there was remarkably little variation among sampled colonies. Some samples did show greater variation across replicates, such as a developing palpon replicate in *Nanomia bijuga* (Fig. S8), and a developing gastrozooid and male gonodendra in *Frillagalma vityazi* (Fig. S7).

The third component of variation we considered was among zooids/specialized tissues within species. This was the greatest component of variation (Fig. S3–S9). We identified genes that significantly differentially expressed in particular zooids, and in the pneumatophore, for each of the sampled species, based on pairwise comparisons (supplementary file 1). In each case, significally differentially expressed refers to genes with higher transcript abundance relative to the other zooid. As it is possible for the same gene to be significantly differentially expressed in pairwise comparisons of several zooids, we also identified a subset of genes that are significantly differentially expressed in only one zooid but not in any other zooid within a given species (Fig. S10, supplementary file 2).

For each species and zooid, we found GO term enrichment for biological processes consistent with the functional specialization of the zooid (supplementary file 3). In gastrozooids that are solely responsible for feeding and digestion, for example, we found these zooids to be enriched for genes involved in chitin, glutathione, and peptide catabolism, proteolysis, as well as metabolism of carbohydrates. Likewise for male gonodendra, we found GO term enrichment for biological processes such as sperm flagellum, mitotic cell cycle process, DNA replication. For female gonodendra, there was GO term enrichment for a number of biological processes including mitotic cell cycle process, DNA replication (in *Agalma elegans*), as well as a number of signalling pathways and developmental processes (in *Frillagalma vityazi*).

Within the four best sampled species, we also identified 92 higher abundance putative transcription factors (24 out of 71 in *Bargmannia elongata*, 43 out of 75 identified in *Frillagalma vityazi*, 50 out of 79 in *Agalma elegans* and 34 out of 72 identified in *Nanomia bijuga*). Many of these transcription factors have higher expression in several zooids regardless of developmental stage (both mature and developing zooids), and a subset have higher expression only in particular zooids and developmental stages (Fig. S11).

### Novel zooids within species: can expression patterns distinguish distinct zooid types?

In siphonophores, there are several instances of lineage-specific zooid diversification events. We investigated gene expression patterns between the novel zooid type and the hypothesized most closely related zooid type in three species. In *Bargmannia elongata* there are two morphologically distinct gastrozooids, that we termed “white” and “yellow” gastrozooids (Fig. S12A and S12B). The “yellow” gastrozooid is larger and darker and occurs as the 7th-10th gastrozooid on the stem [29]. In the Portuguese man of war, *Physalia physalis*, the gastrozooid is unique compared to other gastrozooids in other species - it has a mouth, but no tentacle, and the basigaster region is greatly reduced [9,30]. Meanwhile the tentacle is associated with another zooid, the tentacular palpon (Fig. S12C) [17,30–32]. In *P. physalis*, both the gastrozooid and the tentacular palpon are considered to be subfunctionalized from an ancestral gastrozooid type [31]. Finally, in *Agalma elegans*, there are thought to be at least two different palpon types: gastric palpons that arise at the base of the peduncle of the gastrozooid, and a palpon called the B-palpon (Fig. S12D) [3]. The distinction between these two types of palpon is based on the location of these zooids - the gastrozooid is typically the last element of each cormidium, but based on the budding sequence, Dunn *et al* propose that the enlarged B-palpon is the last element in *A. elegans* [3]. Each of these cases represents a different type of novelty: in *Bargmannia elongata* the distinction between zooids was made based on size and color but not on obvious differences in function, in *P. physalis* the gastrozooids and tentacular palpons differ structurally and functionally, and finally in *A. elegans*, gastric palpons and B palpons differ only in colony location, development, and possibly size.

In *Physalia physalis* 976 genes showed significant differential expression (in all cases, higher transcript abundance relative to the other zooid) in the mature tentacular palpon, compared to 606 genes in the mature gastrozooid (supplementary file 4). A number of genes were significantly differentially expressed in the mature tentacular palpon relative to all other tissues, of which, 670 genes were significantly differentially expressed relative to other tissues that were not shared with the gastrozooid. In the gastrozooid 849 genes were significantly differentially expressed relative to other tissues that were not shared with the tentacular palpon. A number of genes significantly differentially expressed in the tentacular palpon are uncharacterized, however we identified 46 putative toxins in the tentacular palpon, including hemostasis interfering and platelet aggregation activating toxins, phospholipases, serineproteases, hydrolases, metalloendoproteases, calglandulin-like genes, and a neurotoxin (supplementary file 5). By contrast, in the gastrozooid, we found 59 significantly differentially expressed putative toxins (supplementary file 6), including pore-forming Conoporin-Cn1-like and Tereporin-Ca1-like toxins, multiple neurotoxins, hydrolases, serine proteases, toxins likely involved in the promotion of blood coagulation and inhibition of platelet aggregation. Reflecting the role of the gastrozooid in digestion, we also find significant differential expression of digestive enzymes, including chymotrypsin-like genes.

Between the white mature gastrozooid and the yellow mature gastrozooid in *Bargmannia elongata*, few significantly differentially expressed genes were identified (8 genes were up in “white” mature gastrozooids relative to 36 genes up in “yellow” gastrozooids) (supplementary file 4). Among genes that were significantly differentially expressed in either “white” or “yellow” gastrozooids relative to all other tissues, 276 genes were unique to “yellow” gastrozooids and not found in “white” gastrozooids, and 886 genes were found in “white” gastrozooids and not found in “yellow” gastrozooids.

Finally, in *Agalma elegans* very few significantly differentially expressed genes were identified between the B palpon and gastric palpons (1 was significantly differentially expressed in B palpons and 2 were significantly differentially expressed in gastric palpons (supplementary file 4). All three genes have no significant blast hit and did not map to any gene trees). Genes were identified that are significantly differentially expressed in B palpons relative to all other zooids (gastric palpons were excluded). Of this, 928 genes were differentially expressed in the B palpons relative to all other zooids. Most of these genes overlapped with those differentially expressed in the gastric palpons relative to all other zooids (746 genes).

### Classical analysis of gene expression between species: how different is zooid-specific expression between species?

Most comparative studies of gene expression focus exclusively on strict 1:1 orthologs, requiring an assessment of orthology across all species. We call this type of analysis Speciation Tree Filtering (STF), as it limits analyses to a subset of genes with very specific evolutionary histories. Following this approach, we used Orthofinder 2 (v2.4.0) [33] to identify strict orthologs. For all seven species, we identified 1173 strict orthologs, of which 952 orthologs had expression data across all 7 species. In order to increase the number of genes for analysis, we focused on the four best sampled species (*Agalma elegans, Bargmannia elongata, Frillagalma vityazi*, and *Nanomia bijuga*). Using data from 4 taxa, we identified 4009 strict orthologs, of which 3174 orthologs had expression values in four zooids/tissues: gastrozooids (developing and mature), nectophores (developing), palpons (mature), and the pneumatophore. Using TPM10K alone, we found that orthologs clustered largely by species rather than zooid/ tissue (Fig. 3A). Following Breschi et al. [34] (see methods), we used linear models to identify the proportion of variance that could be explained by species or zooid/tissue (Fig. 3B), and found that of 3174 total orthologs, 2125 species-variable genes (SVG), and 168 zooid/tissue-variable genes (TVG).

**Figure 3:**
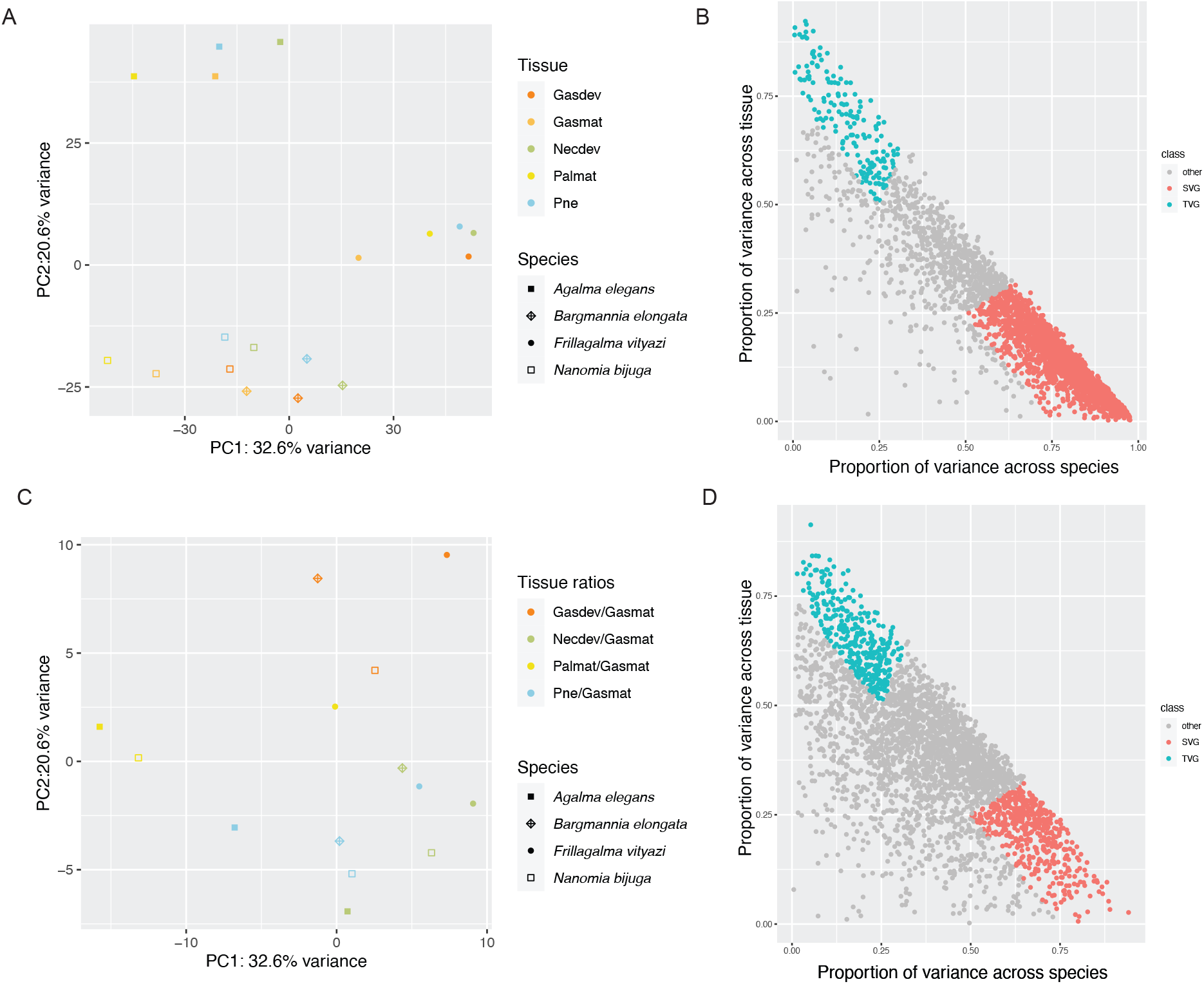
Results of classical gene expression analyses across species, using Speciation Tree Filtering followed by linear models. Top row (A & B): results using TPM10K expression values; Bottom row (C & D): results using TPM10K expression ratios. For both analyses of expression variance explained by species and tissue (right-hand column), we identified highly species-variable genes (SVG, red) and highly tissue-variable genes (TVG, blue). In SVGs, the proportion of variance explained by species is two times greater than that explained by tissues; while in TVGs the proportion of variance explained by tissues is two times greater than that explained by species. All SVGs and TVGs are genes for which both species and tissue explain greater than 75% of variance. A. PCA plot of expression values (TPM10K) of 3174 strict ortholog genes, showing a species dominated clustering pattern. B. The proportion of expression variance explained by species (x-axis) versus the proportion of expression variance explained by tissues (y-axis), using TPM10K expression values. C. PCA plot of expression ratios of 3174 strict ortholog genes, showing a tissue dominated clustering pattern. D. The proportion of expression variance explained by species (x-axis) versus the proportion of expression variance explained by tissues (y-axis), using expression ratios. Abbreviations: Gasdev = developing gastrozooid, Necdev = developing nectophore, Palmat = mature palpon, Pne = pneumatophore, Gasmat = mature gastrozooid.

The strong species-dominated clustering observed when comparing expression values directly may be due to differences in counting efficiency between species, especially as we used reference transcriptomes [19]. To account for this, we used ratios of expression values, using TMP10K expression values of the most commonly sampled zooid – mature gastrozooid – as the denominator [19]. Using this approach, we found that orthologs clustered largely by zooid/tissue rather than species (Fig. 3C). Using linear models, we identified, out of 3174 orthologs, 494 species-variable genes (SVG), and 349 tissue-variable genes (TVG) (Fig. 3D). GO terms enriched among TVG included embryonic morphogenesis, embryo development, cartilage development, and a number of metabolic and biosynthetic processes. Meanwhile, SVG were enriched in GO terms such as cellular response to stress, DNA repair and cell cycle processes. A list of TVG and SVG can be found in supplementary files 7 and 8 respectively.

### Phylogenetic analysis of gene expression with Speciation Branch Filtering: what are the evolutionary changes in zooid expression along branches in the species tree?

For our Speciation Branch Filtering (SBF) analyses (see Fig. 2 and methods), we used transcriptome and genome data from 41 cnidarian species to generate a total of 7070 gene trees, of which 3831 gene trees passed filtering criteria (branch length threshold <2, less than 0.25 branches/tree with default branch lengths, one or more speciation events present in the tree, fewer than 0.3 nodes that have descendant nodes with the same species node ID, that we able to be time calibrated and finally that have a root depth <5). We used Orthofinder 2 [33] to reconcile the species tree and the gene trees and annotate each gene tree node as either a speciation or duplication event. The number of genes represented in these gene trees is shown in Table S1. The internal nodes on these gene trees represent 20088 speciation events and 9082 duplication events. Expression values (TPM10K) were mapped to the tips of the gene trees for each zooid/tissue separately, and ancestral values were inferred at internal nodes with maximum likelihood. We focused on expression in a subset of zooids and tissues that are common across the sampled species: gastrozooids (developing and mature), nectophores (developing), palpons (mature), and the pneumatophore. The distribution of expression changes along species-equivalent branches are shown in figure S13.

As with the STF and linear model analyses, we investigated the impact of counting efficiency on our comparative analyses. We used the TPM10K values at the tips and nodes to calculate expression ratios and calculated changes in expression ratios across a branch for pairs of tissues, using the mature gastrozooid values as the denominator. The distribution of expression ratio changes along species-equivalent branches (branches in the gene tree that correspond to equivalent branches in the species tree, and are descended from speciation events) are shown in figure S14. We found that the variance of change across a given species-equivalent branch is considerably higher for raw TPM10K values as compared with ratios of expression (Figs. S15 & S16). We also identified the number of gene tree branches that had a negative (change <= -2), positive (change >= 2) or neutral (change > -2 and < 2) change (Fig.4). The mean number of species-equivalent branches considered in these analyses are shown in Table 1 – this value reflects the mean number of branches across each zooid or zooid ratio (due to incomplete sampling, some zooids may have fewer or more branches; as seen in Fig.4). For each species-equivalent branch, we found more negative and positive branches than neutral branches when we used TPM10K values, while for expression ratios, we found that the majority of branches show no change across the branch. This suggests that as for the STF analyses, counting efficiency has a large impact on expression change and in turn leads to large gene/branch specific signal. For expression ratios, these results indicate that the vast majority of branches show close to 0 (neutral) change across the branch, suggesting that for closely related genes, expression ratios tend not to differ.

**Figure 4:**
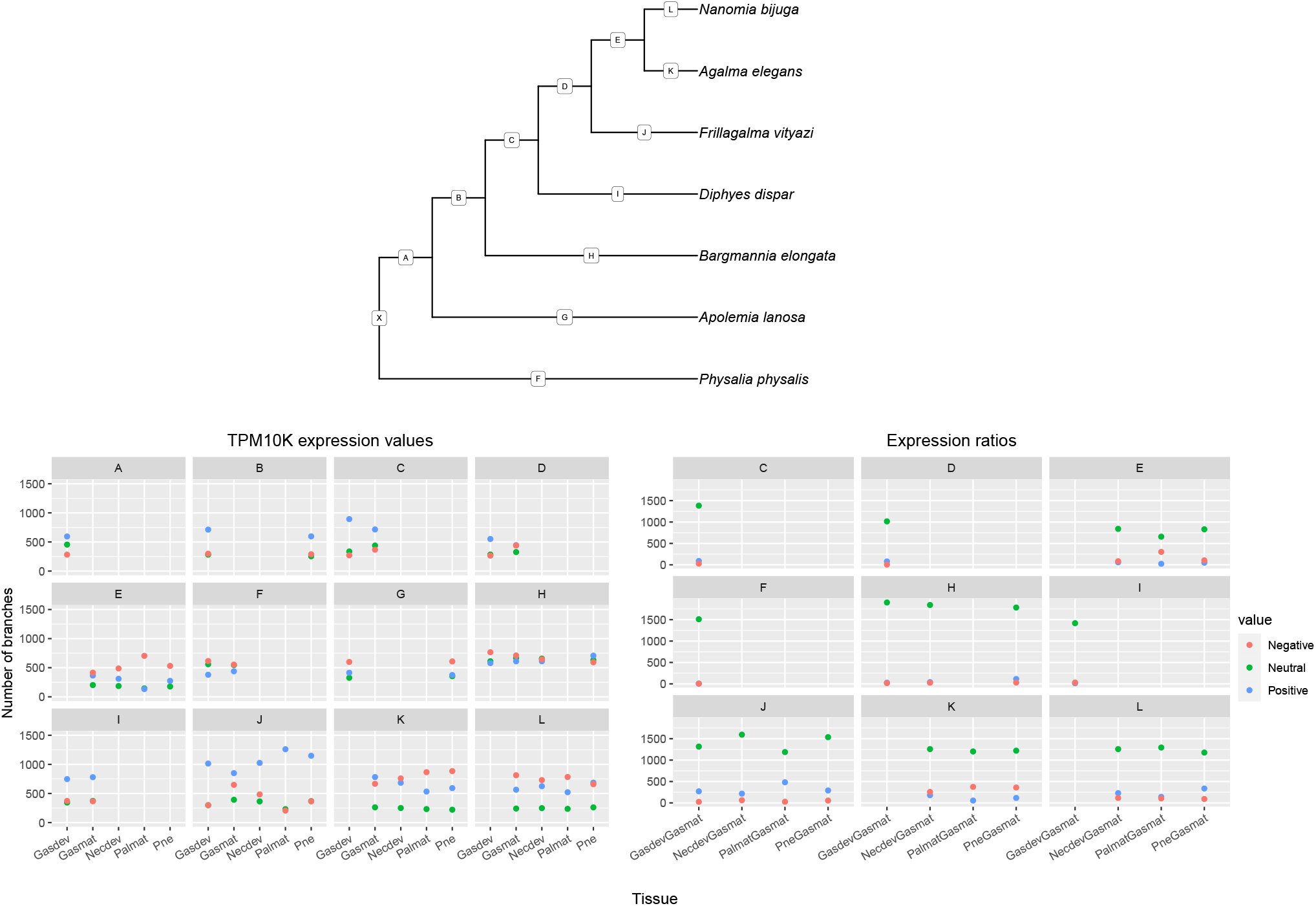
Results of phylogenetic analysis of gene expression analyses, using Speciation Branch Filtering. Number of branches with negative, positive or neutral changes across species-equivalent branches. Top panel: species phylogeny with branch IDs given as letters. Bottom left panel: changes of TPM10K expression across each of the species-equivalent branches (IDs correspond to branch IDs shown in the species tree); bottom right panel: changes of expression ratios across species equivalent branches. Neutral change (green) is defined as <= 2 and >= -2, negative change (red) is defined as <-2, and positive change (blue) is defined as > 2. These units correspond to TPM10K ratios/substitutions per site. GasdevGasmat = developing gastrozooid/ mature gastrozooid, NecdevGasmat = developing nectophore / mature gastrozooid, PalmatGasmat = mature palpon / mature gastrozooid, PneGasmat = Pneumatophore / mature gastrozooid.

**Table 1:** Mean number of species-equivalent branches considered in raw TPM10K and expression ratio analyses (see figure 4 for branch names).

| Species-equivalent branch | Mean Number of Branches TPM10K | Mean Number of Branches ratio |
| --- | --- | --- |
| A | 1331.00 | NA |
| B | 1215.50 | NA |
| C | 1509.50 | 1499.00 |
| D | 1154.50 | 1100.00 |
| E | 982.00 | 982.00 |
| F | 1545.00 | 1521.00 |
| G | 1338.50 | NA |
| H | 1947.00 | 1928.00 |
| I | 1491.00 | 1461.00 |
| J | 1790.60 | 1765.75 |
| K | 1682.75 | 1673.00 |
| L | 1591.75 | 1582.00 |

Out of a total of 3357 final gene trees considered in these analyses, 3329 gene trees contained branches with neutral changes, 1294 gene trees contained branches with positive changes, and 1041 gene trees contained branches with negative changes. Expression ratio changes across species-equivalent branches from all gene trees is available in supplementary file 9 (note: in this file ‘blast hit’ is the most frequent blast hit for the gene tree, gene identity should still be confirmed for each gene, particularly in large gene trees).

The vast majority of gene trees contain species-equivalent branches with neutral changes, including a number of transcription factors and morphogenic signalling pathway genes. For example, in ratios of developing and mature gastrozooids, among the relevant species-equivalent branches in the Wnt gene phylogeny, changes are neutral across Wnt3 and Wnt2 branches, with the exception of a slight positive change across the branch leading to a *Diphyes dispar* Wnt2 paralog, suggesting a higher relative expression of this gene in the developing gastrozooid of *D. dispar* (Fig. S17). For all zooid ratios, we find very consistent Wnt3 expression patterns with neutral or very small positive or negative changes in expression ratio across the branches. Indeed, across all cnidarians examined, Wnt3 has consistent localised expression at the oral pole, likely playing a role in axis formation and maintenance [35–41]. However, for Pneumatophore/Gastrozooid ratios, some large changes are seen across branches K and L, leading to a *Agalma elegans* and *Nanomia bijuga* Wnt2 paralog respectively, likewise for Palpon/Gastrozooid a large change is seen across branch J, leading to a *Frillagalma vityazi* Wnt2 paralog. In *Nematostella vectensis* and *Hydractinia echinata*, Wnt2 is expressed in the middle of the polyp [37,42]. Without more detailed spatial expression patterns, it is difficult to know whether these differences in expression between species reflect species-specific differences in axial patterning and morphogenesis, such as an expansion or reduction of the expression domain.

We also looked at gene trees with species-relative branches that have very large changes (5 < change < -5), representing putative lineage-specific expression patterns. In Frizzled5/8, Wnt4, and homeobox (putatively Hox-B8) gene-trees, for example, we found very large positive changes in ratios of developing gastroozoid/mature gastrozooids across the same species-equivalent branches, J (leading to *Frillagalma vityazi*) and D (leading to Euphysonectae, the clade comprising *A. elegans, N. bijuga*, and *F. vityazi*). In Frizzled5/8 for developing nectophores/gastrozooids we also see a large positive change across branch L (leading to *Nanomia bijuga*) and J (leading to *Frillagalma vityazi*) and negative change across branch K (leading to *Agalma elegans*). In ratios of developing nectophores/gastrozooids, we also found very large positive changes in branch J for Wnt-7b-like, Wnt-4, (putatively) Hox-B8 and NKX1-2. Likewise in pneumatophore/gastrozooid ratios, we also find large positive changes in branch J and L for Frizzled5/8, and a negative change across branch K. Across J, we also see positive changes in pneumatophore/gastrozooid in Hox-B8 and NKX1-2. Finally, for branch K, we see large positive changes in Wnt4 and Wnt7b-like. Many of these genes have been shown to have specific expression domains patterning cnidarian bodies [37,42–44], and further detailed expression analyses of these genes and others within the zooid are required to determine whether the patterns identified here reflect differences in expression domain between species and zooid.

## Discussion

### Evolution of gene expression in Siphonophores

In this study, we used RNA-seq to investigate the functional specialization and evolution of zooids across siphonophore species. In our analysis of differential expression within species, we found a large number of differentially expressed genes across siphonophore zooids, reflecting the distinct anatomy and function of these zooids. In addition, we identified potential transcription factors that are significantly differentially expressed within particular zooids, with potential homologs found in several species, which are interesting candidates for future study in siphonophores or related cnidarian colonial groups. Using these within-species DGE analyses, we identified distinct expression patterns that may be used to define particular zooid types.

We also explored gene expression patterns in three zooid types that are unique to three species, *Physalia physalis, Agalma elegans*, and *Bargmannia elongata*. In siphonophores, different zooid types are typically defined based on morphological and functional differences/similarities, and on the location of the zooid within the colony (which is determined in most species by the asexual budding process in the growth zone, which gives rise to a repeating pattern of zooids along the stem) [3,6,7,29]. Based on both morphology and development, *Physalia physalis* tentacular palpons are considered to be a distinct and unique zooid type [31], and the DGE data indicate that there is a clear morphological and functional difference between gastrozooids and tentacular palpons. We also identified several putative toxin genes with distinct expression profiles between these two zooids, which matches tissue-specific venom observed in other cnidarian species [45–47].

For “yellow” and “white” gastrozooids in *Bargmannia elongata*, the differential expression data point to few morphological or functional differences between these two zooids. Through pairwise differential expression patterns between “yellow” or “white” gastrozooids and other zooids within the colony we were able to identify hundreds of genes that are significantly expressed in one zooid and not the other. This suggests that these two gastrozooids may be distinct zooid types, although they are functionally very similar to one another. Sequencing at greater depth, and also functional work within this species may help clarify the nature of these differences. By contrast, there is no strong evidence in our data that the B palpon and gastric palpons in *Agalma elegans* are sufficiently different to constitute a novel zooid type. These findings suggest that location within the colony is not necessarily sufficient to designate a novel zooid type.

With our STF and linear model analyses we found that the vast majority of identified orthologs were neither exclusively species-variable or tissue-variable, however we were nevertheless able to identify a subset of either tissue- or species-variable genes. As has been observed for vertebrate organs, we find based on GO terms that identified species-variable genes tend to play a role in basic cellular functions (so-called ‘housekeeping genes’), as compared to tissue-variable genes [34].

What can we learn from the SBF analyses, using ratios of expression data? It is important to stress that changes in expression ratios cannot tell us about changes in expression *magnitude*. A tissue-variable gene (identified via linear models in classical analyses) that is either highly or lowly expressed in, for example, nectophores relative to gastrozooids in all species, may show largely neutral changes across all species-equivalent branches. The sign (positive or negative) of change provides an indication of whether expression in the child node is relatively higher in the denominator or the numerator, relative to the parent node. We find overall that most expression ratios show very little change across species-equivalent gene tree branches – suggesting that gene expression patterns tend to be largely consistent among species. Positive or negative changes across branches, especially very large changes, across branches, represent putative linage specific shifts in expression. Whether these reflect distinct morphological or functional changes in the tissues of a particular species or clade requires further validation – a difficulty in this system, where the species are difficult to collect and maintain in the lab.

### Challenges and solutions in the analysis of the evolution of gene expression

In this study we used three different analyses to investigate gene expression patterns within homologous zooids across species, each are complementary and enable different possibilities for biological discovery. Within-species analyses focus on expression among zooids and tissues within a single species – a very large number of genes identified in the transcriptome or genome can be considered, and the broader across species gene-tree need not necessarily be considered. This approach is especially useful for investigating expression patterns within novel zooids that do not have a clear homolog in other species. The classical between species analysis as implemented here, using STF followed by linear models, relies on phylogenetic assumptions yet is a non-phylogenetic tip-focused approach where a single gene is identified per species, and where the identified differences in values across tips are due to changes along all branches in the tree. With linear models, we can identify genes with expression values that are common to a particular zooid across all species, or with expression that is specific to a particular species. However the vast majority of genes are neither zooid or species variable, and show more complex patterns of expression. We proposed a third approach, called Speciation Branch Filtering, that gives access to changes in expression across gene phylogenies, which may differ significantly from the species phylogeny.

Where traditional approaches identify strict ortholog genes *before* conducting analyses, SBF uses information from all gene copies within a gene tree regardless of their evolutionary history of duplication or speciation. By filtering our data by branches, we consider expression patterns at both the tips and internal nodes of gene phylogenies. We are thus able to identify shifts in expression leading to tips as well as shifts in expression leading to particular clades. Although we focus here on expression following speciation events, it is also possible to compare expression following duplication and speciation events.

Using ancestral trait reconstruction, we also overcame sampling issues at the tips of the gene and species phylogeny, as expression values of different treatments can be reconstructed at deep internal nodes, even where there may be inconsistent sampling at the tips. That is, even if expression values are missing for a given tissue and gene in a gene phylogeny, we are still able to examine expression patterns within this gene tree. By contrast, in the Species Tree Filtering analyses, incomplete sampling in the expression matrix leads to the elimination of the ortholog from the analysis.

As with all methods that rely on mapping to reference transcriptomes rather than genomes, this approach is limited by the quality of the reference transcriptomes. Ratios of expression helped significantly to improve issues of differences in count efficiency among species [19]. However, reference transcriptome quality also has an impact on the quality of the gene trees used for expression mapping. Not all reference transcriptomes were sequenced to equal depth among species, and this has important effects on the presence or absence of genes from particular species within the gene tree. This not only has an effect on the representation of expression values, but also impacts the power to investigate patterns of expression among branches within a gene tree. With genome sequencing becoming cheaper and more readily available, the widespread availability of reference genomes will help alleviate many of these issues. Reference genomes will also improve gene models, enabling the distinction of different alleles of the same gene from duplicated gene copies, this in turn will improve the quality of the gene trees. However, gene loss will nevertheless present a challenge to these analyses.

## Conclusions

With the expansion of functional genomic tools, including RNA-seq and single cell sequencing methods, there is considerable interest in looking not only at how genomic variation gives rise to phenotypic diversity in a single species or organism, but also at how functional genomic variation shapes phenotypic diversity across multiple closely and distantly related species to understand broader evolutionary patterns and processes [22–25,34,45,48–57]. Many of these analyses have the goal of identifying shared expression patterns among modular biological units (cell type, tissue, organ, zooid) across species, in order to identify commonalities in expression patterns among species. This is of particular interest for medically orientated fields interested in understanding the extent to which expression results can be extrapolated from one model organism to another. Another goal is to identify expression patterns in a particular biological unit that are unique to a particular species or even clade. For these questions, within-species analyses provide the greatest depth, in terms of number of genes investigated, however comparisons between species on the basis of within-species DGE are limited and largely qualitative. Here we use two approaches to investigate expression patterns in a quantitative manner across homologous tissues. Classic between species analyses, using STF in conjunction with linear models, enabled the investigation of a smaller number of strict orthologs that vary in a tissue or species specific manner, however as this analysis focuses on the tips of gene trees, our ability to investigate lineage or especially clade specific patterns of expression in a given zooid are more limited. Meanwhile phylogenetic analysis using SBF focuses on branches rather than tips, also with specific evolutionary histories (descended from speciation events), but it enables the identification of expression patterns that vary little among genes/species, as well as expression patterns that show strong lineage or clade specific patterns for a given zooid.

## Methods

All scripts for the analyses are available in a git repository at https://github.com/dunnlab/Siphonophore_Expression. Additional data files required for analyses that are too large for the github repository can be found on Figshare with the following DOIs: 10.6084/m9.figshare.14838384, 10.6084/m9.figshare.14838372, 10.6084/m9.figshare.14838315, 10.6084/m9.figshare.14838090, 10.6084/m9.figshare.14829183. The most recent commit at the time of the analysis presented here was fc063e9.

### Collecting

Specimens were collected in the north-eastern Pacific Ocean in Monterey Bay and, in the case of *Physalia physalis* the Gulf of Mexico. Specimens were collected by remotely operated vehicle (ROV) or during blue-water SCUBA dives. *Physalia* specimens were collected by hand from the beach after being freshly washed on-shore by prevailing winds. Available physical vouchers have been deposited at the Peabody Museum of Natural History (Yale University), New Haven, CT. Specimens were relaxed using 7.5% MgCl_2_ hexahydrate in Milli-Q water at a ratio of 1/3 MgCl_2_ and 2/3 seawater. Zooids were subsequently dissected from the colony and flash frozen in liquid nitrogen. Colonies were cooled to collection temperatures (e.g 4 degrees C for deep sea species) while the dissections took place. Dissections took no longer than 15-20 minutes. In the case of large colonies, the stem was cut and only partial sections of the colony were placed under the microscope at a given time. Each replicate individual represents a genetically distinct colony from the same species. Replicate specimens were of an equivalent colony size, and zooid replicates were also equivalent sizes. Larger zooid types, such as gastrozooids, were sampled as a single zooid, but smaller zooids were pooled. Pooled zooids were of a comparable maturity and sampled from the same location in a single colony. Sampling data, including time, date, depth, and voucher ID, can be found in supplementary file 10.

### Sequencing

mRNA was extracted directly from tissue using Zymo Quick RNA MicroPrep (Zymo #R1050), including a DNase step, and subsequently prepared for sequencing using the Illumina TruSeq Stranded Library Prep Kit (Illumina, #RS-122-2101). 50 base-pair single-end libraries were all sequenced on the HiSeq 2500 sequencing platform. Three sequencing runs were conducted, representing three full flow cells. To avoid potential run/lane confounding effects, where possible, libraries of multiple zooids/tissue of a single individual in a species were barcoded and pooled in a single sequencing lane, and replicate lanes of zooids/tissue from different individuals of the same species were sequenced in separate runs. Additionally, two libraries were run as technical replicates across all runs and many lanes, for a total of 20 technical replicates. Sequences, along with sampling metadata, are publicly available on NCBI under BioProject ID PRJNA540747.

### Analysis

#### Differential gene expression

Short read libraries were mapped to previously published transcriptomes [26] using Agalma v 2.0.0 [58,59], which uses a number of existing tools for transcript quantification, including RSEM (which uses Bowtie) [20,60]. Using the `agalmar` package (https://github.com/caseywdunn/agalmar), we filtered out genes that were flagged as being rRNA, and selected only protein coding genes. We also only considered genes that were greater than 0 in at least two libraries. Differential gene expression analyses, including normalization, were conducted in R, using the `DESeq2` package [61]. Libraries that were found to be outliers based on mean Cook’s distance were removed from the DESeq object and from downstream analyses and normalization. Testing for differential expression was conducted using the `results()` function in DESeq2. Genes were considered to be significantly differentially expressed if adjusted p-values (Bonferroni correction) were less than 0.05. Differential expression analyses were only conducted on zooids/tissue with two or more replicates.

GO annotations were retrieved for each of the reference translated transcriptomes [26] using the PANNZER2 web server [62].The PANNZER2 format was modified to match the gene2GO format required for the package `topGO` [63]. Gene set enrichment analyses were carried out within species using the R package `GOseq` [64], which takes gene length into account. Over and underrepresented categories were calculated using the Wallenius approximation, and p-values were adjusted using the Benjamini and Hochberg method. Categories with an adjusted p-value below 0.05 are considered enriched. Gene set enrichment analyses were also conducted at the gene tree level, considering representative GO terms for particular gene trees. Representative GO terms were selected based on the frequency of occurrence among genes in the gene tree. As gene lengths vary among species and genes in the gene tree, the `GOseq` approach could not be used, and `topGO` was used to detect GO terms that are enriched based on Fisher’s exact test. This approach assumes that each gene tree has an equal probability of having genes shared among species that are detected as differentially expressed, however results may be biased by a number of factors, including mean gene length among genes in the gene tree [64]. Putative toxin genes were identified using blastp from the ToxProt dataset (http://www.uniprot.org/program/Toxins, last accessed 13 May 2021). See Supplementary Information for R package version numbers.

#### TPM10K

For all comparative expression analyses, expression values were normalized using a new method we call transcripts per million 10K (TPM10K). For gene *i* of a given species, TPM is typically calculated as [65]:

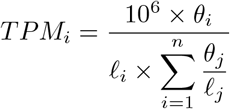

Where *θ*_*i*_ is the number of the mapped reads to gene *i, l*_*i*_ is the effective length of the gene, and n is the number of genes in the reference. The intent of this measure is to make libraries comparable within a single species. The sum of TPM values within a library is 10^6^, and the mean is 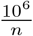. One implication of this is that TPM values are not directly comparable across species, since in practice *n* differs across species. If this were not accounted for, then it could appear, for example, that genes all have lower expression in a species with a more complete reference transcriptome and higher *n*. To account for differences in means among species, we use a new measure, TPM10K, that accounts for differences in *n*:

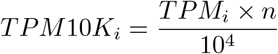

Where the sum of TPM10K values within a library is 10^2^ × *n* and the mean is 10^2^. By multiplying by *n* we are able to account for different sequencing depths among species, and ensure a common mean. As *n* is large, we divide by an arbitrary number (in this case 10^4^) in order to reduce the magnitude of the expression value.

#### Speciation Tree Filtering

For Speciation Tree Filtering (STF) analyses, single copy orthologs were obtained using Orthofinder 2 (v2.4.0) [33]. Linear models used in STF analyses were constructed using `lm()`, following methods and code developed by Breschi et al. 2016 [34], https://github.com/abreschi/Rscripts/blob/master/anova.R. All SVGs and TVGs are genes for which both species and tissue explain greater than 75% of variance. Additionally, in SVGs, the proportion of variance explained by species is two times greater than that explained by tissues; while in TVGs the proportion of variance explained by tissues is two times greater than that explained by species.

#### Speciation Branch Filtering

Gene trees were generated using the transcriptomes and genomes from 41 species [26] using Agalma v 2.0.0 [58,59]. Following the `treefinder` step of Agalma v 2.0.0, amino acid data were exported and supplied to Orthofinder 2 (v2.3.8) [33] for simultaneous co-estimation of gene trees with the published maximum likelihood species tree [26]. Within Orthofinder 2, the selected multiple sequence alignment method was MAFFT [66] and maximum likelihood tree inference method was IQ-tree with the LG+F+R4 substitution model [67].

Phylogenetic analyses were conducted in `R` using `geiger`, `ape`, `phytools`, `Rphylopars`, and `hutan` [68–72]. Phylogenetic trees were visualized in `R` using `ggtree` and `treeio` [73].

Gene trees were filtered to exclude trees with a length threshold > 2 and that had more than 0.25 branches with a default length value (that are indicative of branch length=0). Gene trees nodes were annotated as speciation events or duplication events, based on assignment by Orthofinder 2 [33]. Speciation nodes were subsequently assigned a node ID equivalent to the species tree node, using species tip names from the gene tree to determine the most common recent ancestor in the species tree, using the phytools package. Due to the use of the species-overlap method by Orthofinder 2, some clades of single copy genes were assigned as speciation events, although the topology is inconsistent with the species tree. To avoid time calibration issues, due to descendant nodes being assigned the same species node ID, descendant speciation nodes with the same species node ID were marked as null, and gene trees with nodes greater than 0.3 null nodes per internal node were excluded – indicating widespread topological differences with the species tree. Additionally, only trees with one or more speciation events were retained, as speciation events are used for time calibrations. The gene trees were then time calibrated to the species tree using `chronos()` in the ape package, so that the branch lengths were scaled to the same equivalent length across all gene trees [68]. Some gene trees could not be calibrated against the node constraints from the species tree and were discarded, additionally gene trees with roots that exceeded a maximum root depth of 5 were excluded. Tips without expression values were then pruned out of the tree. Gene trees with fewer than three expression values at the tips were discarded, retaining only trees with three or more values. Pruned and unpruned calibrated gene tree files are available as supplementary data.

We then took the mean TPM10K value for each gene across replicates of the same zooid/tissue within a species and applied a log transformation. Using gene trees with expression values for each gene within a species at the tips, maximum likelihood ancestral trait values were generated at the nodes using the anc.recon() function in `Rphylopars` assuming a Brownian Motion model of evolution (Fig. 2, step 2 & 3) [70]. As not all zooids are present in all of the species, the trees were pruned down to the subset of tips with expression values for ancestral trait reconstructions. Node values were then added back to the unpruned tree with all of the reconstructed expression values. Change in expression was measured across a branch by taking the difference between a parent node and a child node, and then this difference is scaled by branch length (Fig. 2, step 4).

To examine if expression values in our dataset evolve under Brownian Motion (BM), we simulated data set of random expression values using `fastBM()` from the `phytools` package with empirically derived mean and standard-deviation values for each gene tree. Under this model, expression variance accumulates among linages on the gene tree as a function of time, and is used as a model of evolution under drift, as well as some forms of natural selection [74–76].

## Supporting information

Gene trees

## Acknowledgements

This work was supported by the National Science Foundation (DEB-1256695, the Waterman Award). CM was also supported in part by a RI-EPSCoR Fellowship, NSF EPS-1004057. Analyses were conducted with computational resources and services at the Center for Computation and Visualization at Brown University, supported in part by the NSF EPSCoR EPS-1004057 and the State of Rhode Island. Analyses were also conducted at the Yale Center for Research Computing (YCRC). We would like to thank Steve Haddock, Lynne Christianson, and the whole crew of the Western Flyer for their help and expertise. Additional thanks are due to Samuel Church for his assistance during a research cruise, and for productive conversations about comparative gene expression methods.

## References

1. Hiebert LS, Simpson C, Tiozzo S: Coloniality, clonality, and modularity in animals: The elephant in the room. Journal of Experimental Zoology Part B: Molecular and Developmental Evolution 2020, 336:198–211, https://doi.org/10.1002/jez.b.22944.

2. Beklemishev WN: Principles of Comparative Anatomy of Invertebrates. Volume I. Promorphology. Edinburgh: Oliver & Boyd; 1969.

3. Dunn CW, Wagner GP: The evolution of colony-level development in the Siphonophora (Cnidaria: Hydrozoa). Development Genes and Evolution 2006, 216:743–754, https://doi.org/10.1007/s00427-006-0101-8.

4. Mackie GO: Siphonophores, Bud Colonies, and Superorganism. In The lower metazoa. Edited by Dougherty E Berkeley: University of California Press; 1963:329–337.

5. Mackie GO: From aggregates to integrates: physiological aspects of modularity in colonial animals. Philosophical Transactions of the Royal Society of London. Series B, Biological sciences 1986, 313:175–196.

6. Mackie GO, Pugh PR, Purcell JE: Siphonophore biology. Advances in Marine Biology 1987, 24:97–262, https://doi.org/10.1016/S0065-2881(08)60074-7.

7. Totton AK: A synopsis of the Siphonophora. British Museum (Natural History); 1965.

8. Church SH, Siebert S, Bhattacharyya P, Dunn CW: The histology of Nanomia bijuga (Hydrozoa: Siphonophora). Journal of Experimental Zoology Part B: Molecular and Developmental Evolution 2015, 324:435–449, https://doi.org/10.1002/jez.b.22629.

9. Mackie GO: Studies on Physalia physalis (L.). Part 2. Behavior and histology. Discovery Reports 1960, 30:371407.

10. Carré C: Le developpement larvaire d’Abylopsis tetragona. Cahiers de Biologie Marine 1967, 8:185–193.

11. Carré D: Etude histologique du developpement de Nanomia bijuga (Chiaje, 1841), Siphonophore Physonecte, Agalmidae. Cahiers de Biologie Marine 1969, 10:325–341.

12. Carré C, Carré D: A complete life cycle of the calycophoran siphonophore Muggiaea kochi (Will) in the laboratory, under different temperature conditions: ecological implications. Philosophical Transactions of the Royal Society of London. Series B, Biological sciences 1991, 334:27–32, https://doi.org/10.1098/rstb.1991.0095.

13. Siebert S, Goetz FE, Church SH, Bhattacharyya P, Zapata F, Haddock SHD, Dunn CW: Stem cells in Nanomia bijuga (Siphonophora), a colonial animal with localized growth zones. EvoDevo 2015, 6:22, https://doi.org/10.1186/s13227-015-0018-2.

14. Sanders SM, Shcheglovitova M, Cartwright P: Differential gene expression between functionally specialized polyps of the colonial hydrozoan Hydractinia symbiolongicarpus (Phylum Cnidaria). BMC Genomics 2014, 15:406, https://doi.org/10.1093/gbe/evv153.

15. Sanders SM, Cartwright P: Interspecific differential expression analysis of RNA-Seq data yields insight into life cycle variation in hydractiniid hydrozoans. Genome Biology and Evolution 2015, 7:2417–2431, https://doi.org/10.1186/1471-2164-15-406.

16. Siebert S, Robinson MD, Tintori SC, Goetz F, Helm RR, Smith SA, Shaner N, Haddock SHD, Dunn CW: Differential Gene Expression in the Siphonophore Nanomia bijuga (Cnidaria) Assessed with Multiple Next-Generation Sequencing Workflows. PLoS ONE 2011, 6:e22953, https://doi.org/10.1371/journal.pone.0022953.g007.

17. Bardi J, Marques AC: Taxonomic redescription of the Portuguese man-of-war, Physalia physalis (Cnidaria, Hydrozoa, Siphonophorae, Cystonectae) from Brazil. Iheringia, Série Zoologia 2007, 97:425–433, https://doi.org/10.1590/S0073-47212007000400011.

18. Macrander J, Brugler MR, Daly M: A RNA-seq approach to identify putative toxins from acrorhagi in aggressive and non-aggressive Anthopleura elegantissima polyps. BMC Genomics 2015, 16:221, https://doi.org/10.1186/s12864-015-1417-4.

19. Dunn CW, Luo X, Wu Z: Phylogenetic analysis of gene expression. Integrative and Comparative Biology 2013, 53:847–856, https://doi.org/10.1093/icb/ict068.

20. Li B, Dewey CN: RSEM: accurate transcript quantification from RNA-Seq data with or without a reference genome. BMC Bioinformatics 2011, 12:323, https://doi.org/10.1186/1471-2105-12-323.

21. Wagner GP, Kin K, Lynch VJ: Measurement of mRNA abundance using rna-seq data: RPKM measure is inconsistent among samples. Theory in biosciences 2012, 131:281–285, https://doi.org/10.1007/s12064-012-0162-3.

22. Brawand D, Soumillon M, Necsulea A, Julien P, Csárdi G, Harrigan P, Weier M, Liechti A, Aximu-Petri A, Kircher M, Albert FW, Zeller U, Khaitovich P, Grützner F, Bergmann S, Nielsen R, Pääbo S, Kaessmann H: The evolution of gene expression levels in mammalian organs. Nature 2011, 478:343–348, https://doi.org/10.1038/nature10532.

23. Cardoso-Moreira M, Halbert J, Valloton D, Velten B, Chen C, Shao Y, Liechti A, Ascenção K, Rummel C, Ovchinnikova S, Mazin PV, Xenarios I, Harshman K, Mort M, Cooper DN, Sandi C, Soares MJ, Ferreira PG, Afonso S, Carneiro M, Turner JMA, VandeBerg JL, Fallahshahroudi A, Jensen P, Behr R, Lisgo S, Lindsay S, Khaitovich P, Huber W, Baker J, et al.: Gene expression across mammalian organ development. Nature 2019, 571:505–509, https://doi.org/10.1038/s41586-019-1338-5.

24. Levin M, Anavy L, Cole AG, Winter E, Mostov N, Khair S, Senderovich N, Kovalev E, Silver DH, Feder M, Fernandez-Valverde SL, Nakanishi N, Simmons D, Simakov O, Larsson T, Liu S-Y, Jerafi-Vider A, Yaniv K, Ryan JF, Martindale MQ, Rink JC, Arendt D, Degnan SM, Degnan BM, Hashimshony T, Yanai I: The mid-developmental transition and the evolution of animal body plans. Nature 2016, 531:637–641, https://doi.org/10.1038/nature16994.

25. Yang R, Wang X: Organ evolution in angiosperms driven by correlated divergences of gene sequences and expression patterns. The Plant Cell 2013, 25:71–82, https://doi.org/10.1105/tpc.112.106716.

26. Munro C, Siebert S, Zapata F, Howison M, Damian-Serrano A, Church SH, Goetz FE, Pugh PR, Haddock SHD, Dunn CW: Improved phylogenetic resolution within Siphonophora (Cnidaria) with implications for trait evolution. Molecular Phylogenetics and Evolution 2018, 127:823–833, https://doi.org/10.1016/j.ympev.2018.06.030.

27. Auer PL, Doerge R: Statistical design and analysis of RNA-Seq data. Genetics 2010, 185:405–416, https://doi.org/10.1534/genetics.110.114983.

28. McIntyre LM, Lopiano KK, Morse AM, Amin V, Oberg AL, Young LJ, Nuzhdin SV: RNA-seq: technical variability and sampling. BMC Genomics 2011, 12:293, https://doi.org/10.1186/1471-2164-12-293.

29. Dunn CW: Complex colony-level organization of the deep-sea siphonophore Bargmannia elongata (Cnidaria, Hydrozoa) is directionally asymmetric and arises by the subdivision of pro-buds. Developmental Dynamics 2005, 234:835–845, https://doi.org/10.1002/dvdy.20483.

30. Totton AK: Studies on Physalia physalis (L.). Part 1. Natural history and morphology. Discovery Reports 1960, 30:301–368.

31. Munro C, Vue Z, Behringer RR, Dunn CW: Morphology and development of the Portuguese man of war, Physalia physalis. Scientific reports 2019, 9:15522, https://doi.org/10.1038/s41598-019-51842-1.

32. Haeckel E: Report on the Siphonophorae collected by HMS Challenger during the years 1873-1876. Report of the Scientific Results of the voyage of H.M.S. Challenger. Zoology 1888, 28:1–380.

33. Emms DM, Kelly S: OrthoFinder: Phylogenetic orthology inference for comparative genomics. Genome biology 2019, 20:238, https://doi.org/10.1186/s13059-019-1832-y.

34. Breschi A, Djebali S, Gillis J, Pervouchine DD, Dobin A, Davis CA, Gingeras TR, Guigó R: Genespecific patterns of expression variation across organs and species. Genome Biology 2016, 17:151, https://doi.org/10.1186/s13059-016-1008-y.

35. Bagaeva TS, Kupaeva DM, Vetrova AA, Kosevich IA, Kraus YA, Kremnyov SV: cWnt signaling modulation results in a change of the colony architecture in a hydrozoan. Developmental biology 2019, 456:145–153, https://doi.org/10.1016/j.ydbio.2019.08.019.

36. Guder C, Philipp I, Lengfeld T, Watanabe H, Hobmayer B, Holstein T: The wnt code: Cnidarians signal the way. Oncogene 2006, 25:7450–7460, https://doi.org/10.1038/sj.onc.1210052.

37. Hensel K, Lotan T, Sanders SM, Cartwright P, Frank U: Lineage-specific evolution of cnidarian Wnt ligands. Evolution & Development 2014, 16:259–269, https://doi.org/10.1111/ede.12089.

38. Hobmayer B, Rentzsch F, Kuhn K, Happel CM, Laue CC von, Snyder P, Rothbächer U, Holstein TW: WNT signalling molecules act in axis formation in the diploblastic metazoan Hydra. Nature 2000, 407:186–189, https://doi.org/10.1038/35025063.

39. Momose T, Derelle R, Houliston E: A maternally localised Wnt ligand required for axial patterning in the cnidarian Clytia hemisphaerica. Development 2008, 135:2105–2113.

40. Nawrocki AM, Cartwright P: Expression of Wnt pathway genes in polyps and medusa-like structures of Ectopleura larynx (Cnidaria: Hydrozoa). Evolution & development 2013, 15:373–384, https://doi.org/10.1111/ede.12045.

41. Sanders SM, Cartwright P: Patterns of Wnt signaling in the life cycle of Podocoryna carnea and its implications for medusae evolution in Hydrozoa (Cnidaria). Evolution & development 2015, 17:325–336, https://doi.org/10.1111/ede.12165.

42. Kusserow A, Pang K, Sturm C, Hrouda M, Lentfer J, Schmidt HA, Technau U, Von Haeseler A, Hobmayer B, Martindale MQ, Holstein TW: Unexpected complexity of the Wnt gene family in a sea anemone. Nature 2005, 433:156–160, https://doi.org/10.1038/nature03158.

43. Ryan JF, Mazza ME, Pang K, Matus DQ, Baxevanis AD, Martindale MQ, Finnerty JR: Pre-bilaterian origins of the Hox cluster and the Hox code: evidence from the sea anemone, Nematostella vectensis. PloS one 2007, 2:e153, https://doi.org/10.1371/journal.pone.0000153.

44. Sinigaglia C, Busengdal H, Leclere L, Technau U, Rentzsch F: The bilaterian head patterning gene six3/6 controls aboral domain development in a cnidarian. PLoS Biol 2013, 11:e1001488, https://doi.org/10.1371/journal.pbio.1001488.

45. Macrander J, Broe M, Daly M: Tissue-Specific Venom Composition and Differential Gene Expression in Sea Anemones. Genome Biology and Evolution 2016, 8:2358–2375, https://doi.org/10.1093/gbe/evw155.

46. Ames CL, Ryan JF, Bely AE, Cartwright P, Collins AG: A new transcriptome and transcriptome profiling of adult and larval tissue in the box jellyfish Alatina alata: an emerging model for studying venom, vision and sex. BMC genomics 2016, 17:650, https://doi.org/10.1186/s12864-016-2944-3.

47. Klompen AML, Kayal E, Collins AG, Cartwright P: Phylogenetic and selection analysis of an expanded family of putatively pore-forming jellyfish toxins (Cnidaria: Medusozoa). Genome Biology and Evolution 2021, 13:evab081, https://doi.org/10.1093/gbe/evab081.

48. Barbosa-Morais NL, Irimia M, Pan Q, Xiong HY, Gueroussov S, Lee LJ, Slobodeniuc V, Kutter C, Watt S, Colak R, others: The evolutionary landscape of alternative splicing in vertebrate species. Science 2012, 338:1587–1593, https://doi.org/10.1126/science.1230612.

49. Clarke TH, Garb JE, Haney RA, Chaw RC, Hayashi CY, Ayoub NA: Evolutionary shifts in gene expression decoupled from gene duplication across functionally distinct spider silk glands. Scientific Reports 2017, 7:8393, https://doi.org/10.1038/s41598-017-07388-1.

50. Merkin J, Russell C, Chen P, Burge CB: Evolutionary dynamics of gene and isoform regulation in Mammalian tissues. Science 2012, 338:1593–1599, https://doi.org/10.1126/science.1228186.

51. Necsulea A, Soumillon M, Warnefors M, Liechti A, Daish T, Zeller U, Baker JC, Grützner F, Kaessmann H: The evolution of lncRNA repertoires and expression patterns in tetrapods. Nature 2014, 505:635–640, https://doi.org/10.1038/nature12943.

52. Perry GH, Melsted P, Marioni JC, Wang Y, Bainer R, Pickrell JK, Michelini K, Zehr S, Yoder AD, Stephens M, Pritchard JK, Yoav G: Comparative RNA sequencing reveals substantial genetic variation in endangered primates. Genome Research 2012, 22:602–610, https://doi.org/10.1101/gr.130468.111.

53. Sudmant PH, Alexis MS, Burge CB: Meta-analysis of RNA-seq expression data across species, tissues and studies. Genome Biology 2015, 16:287, https://doi.org/10.1186/s13059-015-0853-4.

54. Ma S, Avanesov AS, Porter E, Lee BC, Mariotti M, Zemskaya N, Guigo R, Moskalev AA, Gladyshev VN: Comparative transcriptomics across 14 Drosophila species reveals signatures of longevity. Aging Cell 2018, 17:e12740.

55. Zhang G, Li C, Li Q, Li B, Larkin DM, Lee C, Storz JF, Antunes A, Greenwold MJ, Meredith RW, Ödeen A, Cui J, Zhou Q, Xu L, Pan H, Wang Z, Jin L, Zhang P, Hu H, Yang W, Hu J, Xiao J, Yang Z, Liu Y, Xie Q, Yu H, Lian J, Wen P, Zhang F, Li H, et al.: Comparative genomics reveals insights into avian genome evolution and adaptation. Science 2014, 346:1311–1320, https://doi.org/10.1126/science.1251385.

56. Darbellay F, Necsulea A: Comparative Transcriptomics Analyses across Species, Organs, and Developmental Stages Reveal Functionally Constrained lncRNAs. Molecular Biology and Evolution 2019, 37:240–259, https://doi.org/10.1093/molbev/msz212.

57. Fukushima K, Pollock DD: Amalgamated cross-species transcriptomes reveal organ-specific propensity in gene expression evolution. Nature communications 2020, 11:4459, https://doi.org/10.1038/s41467-020-18090-8.

58. Dunn CW, Howison M, Zapata F: Agalma: an automated phylogenomics workflow. BMC Bioinformatics 2013, 14:330, https://doi.org/10.1186/1471-2105-14-330.

59. Guang A, Howison M, Zapata F, Lawrence C, Dunn CW: Revising transcriptome assemblies with phylogenetic information. PLoS One 2021, 16:e0244202, https://doi.org/10.1371/journal.pone.0244202.

60. Langmead B, Trapnell C, Pop M, Salzberg SL: Ultrafast and memory-efficient alignment of short DNA sequences to the human genome. Genome Biology 2009, 10:R25, https://doi.org/10.1186/gb-2009-10-3-r25.

61. Love MI, Huber W, Anders S: Moderated estimation of fold change and dispersion for RNA-seq data with DESeq2. Genome Biology 2014, 15:550, https://doi.org/10.1186/s13059-014-0550-8.

62. Törönen P, Medlar A, Holm L: PANNZER2: a rapid functional annotation web server. Nucleic Acids Research 2018, 46:W84–W88, https://doi.org/10.1093/nar/gky350.

63. Alexa A, Rahnenfuhrer J: TopGO: Enrichment analysis for gene ontology. R package version 2.40.0; 2020.

64. Young MD, Wakefield MJ, Smyth GK, Oshlack A: Gene ontology analysis for RNA-seq: accounting for selection bias. Genome Biology 2010, 11:R14, https://doi.org/10.1186/gb-2010-11-2-r14.

65. Li B, Ruotti V, Stewart RM, Thomson JA, Dewey CN: RNA-Seq gene expression estimation with read mapping uncertainty. Bioinformatics 2009, 26:493–500, https://doi.org/10.1093/bioinformatics/btp692.

66. Katoh K, Standley DM: MAFFT multiple sequence alignment software version 7: Improvements in performance and usability. Molecular biology and evolution 2013, 30:772–780.

67. Nguyen L-T, Schmidt HA, Von Haeseler A, Minh BQ: IQ-tree: A fast and effective stochastic algorithm for estimating maximum-likelihood phylogenies. Molecular biology and evolution 2015, 32:268–274, https://doi.org/10.1093/molbev/msu300.

68. Paradis E, Claude J, Strimmer K: APE: analyses of phylogenetics and evolution in R language. Bioinformatics 2004, 20:289–290, https://doi.org/10.1093/bioinformatics/btg412.

69. Church SH, Ryan JF, Dunn CW: Automation and Evaluation of the SOWH Test with SOWHAT. Systematic Biology 2015, 64:1048–105810.1093/sysbio/syv055.

70. Goolsby EW, Bruggeman J, Ané C: Rphylopars: fast multivariate phylogenetic comparative methods for missing data and within-species variation. Methods in Ecology and Evolution 2017, 8:22–27, https://doi.org/10.1111/2041-210X.12612.

71. Harmon LJ, Weir JT, Brock CD, Glor RE, Challenger W: GEIGER: investigating evolutionary radiations. Bioinformatics 2007, 24:129–131, https://doi.org/10.1093/bioinformatics/btm538.

72. Revell LJ: phytools: an R package for phylogenetic comparative biology (and other things). Methods in Ecology and Evolution 2012, 3:217–223, https://doi.org/10.1111/j.2041-210X.2011.00169.x.

73. Yu G, Smith DK, Zhu H, Guan Y, Lam TT-Y: ggtree: an R package for visualization and annotation of phylogenetic trees with their covariates and other associated data. Methods in Ecology and Evolution 2017, 8:28–36, https://doi.org/10.1111/2041-210X.12628.

74. Felsenstein J: Maximum-likelihood estimation of evolutionary trees from continuous characters. American journal of human genetics 1973, 25:471.

75. Revell LJ, Harmon LJ: Testing quantitative genetic hypotheses about the evolutionary rate matrix for continuous characters. Evolutionary Ecology Research 2008, 10:311–331.

76. Revell LJ, Harmon LJ, Collar DC: Phylogenetic signal, evolutionary process, and rate. Systematic biology 2008, 57:591–601, https://doi.org/10.1080/10635150802302427.

