## Supplementary figures and tables for "Evolution of gene expression across species and specialized zooids in Siphonophora"

### Supplementary data

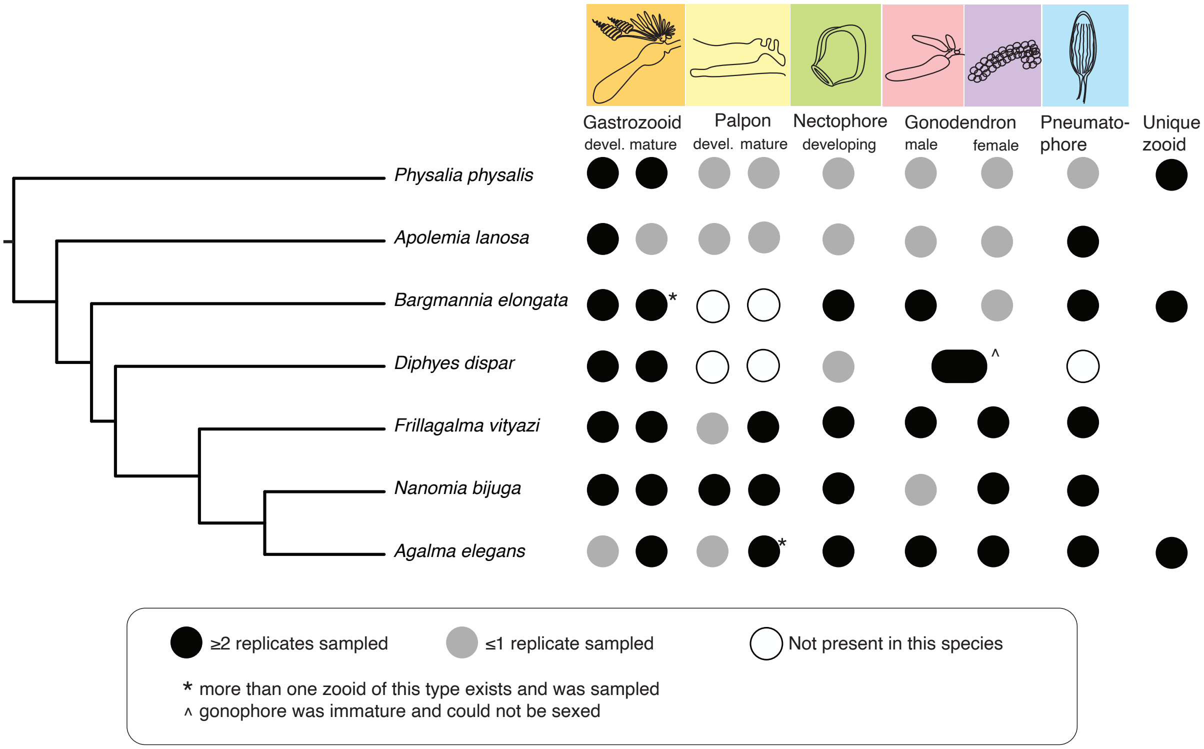


Figure S1: Phylogeny of the focal species sampled in this study, with details of the traits sampled for each of the species. Phylogeny modified from Munro et al. 2018, [26]. Black indicates that multiple replicates have been sampled, grey indicates that no or only one replicate has been sampled, and white indicates that this zooid/tissue is not present in this species. The category "unique zooid" indicates that a zooid type that is unique to this species was sampled.

Table S1: Number of genes per species (library size), number of sampled zooids (at least two replicates each, including developing and mature stages), total number of genes in gene trees (there can be multiple genes in a gene tree), and number of unique gene trees containing genes from this species.

| Species | Number of Genes | Number of Zooids Sampled | Number of Genes in Gene Trees | Total Number of Gene Trees |
|---|---|---|---|---|
| *Diphyes dispar* | 51981 | 3 | 5833 | 3381 |
| *Frillagalma vityazi* | 49145 | 8 | 5186 | 3394 |
| *Nanomia bijuga* | 36249 | 7 | 5053 | 3274 |
| *Agalma elegans* | 30234 | 7 | 4962 | 3488 |
| *Bargmannia elongata* | 38177 | 7 | 4550 | 3401 |
| *Physalia physalis* | 19965 | 5 | 3886 | 2789 |
| *Apolemia lanosa* | 18541 | 2 | 3531 | 2628 |

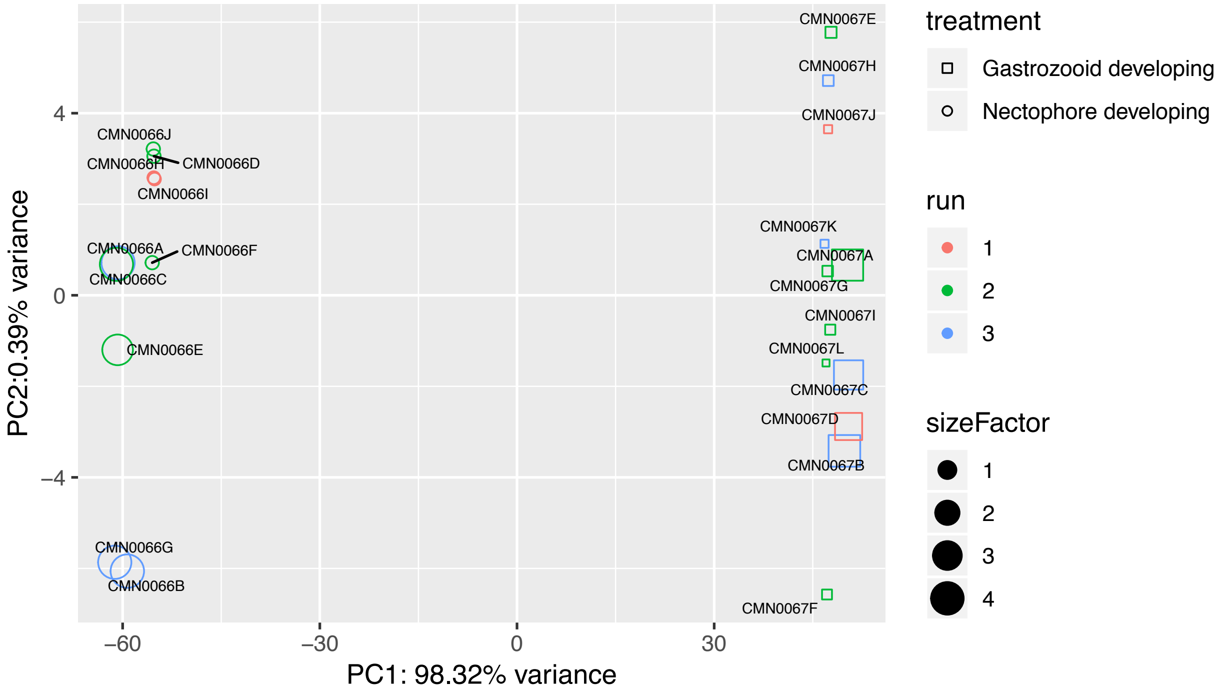


Figure S2: PCA of regularized log transformed expression counts of technical replicates of two different zooids in *Frillagalma vityazi* from different runs and lanes. Color indicates the run number, shape indicates the zooid, and size factor indicates number of genes sequenced.

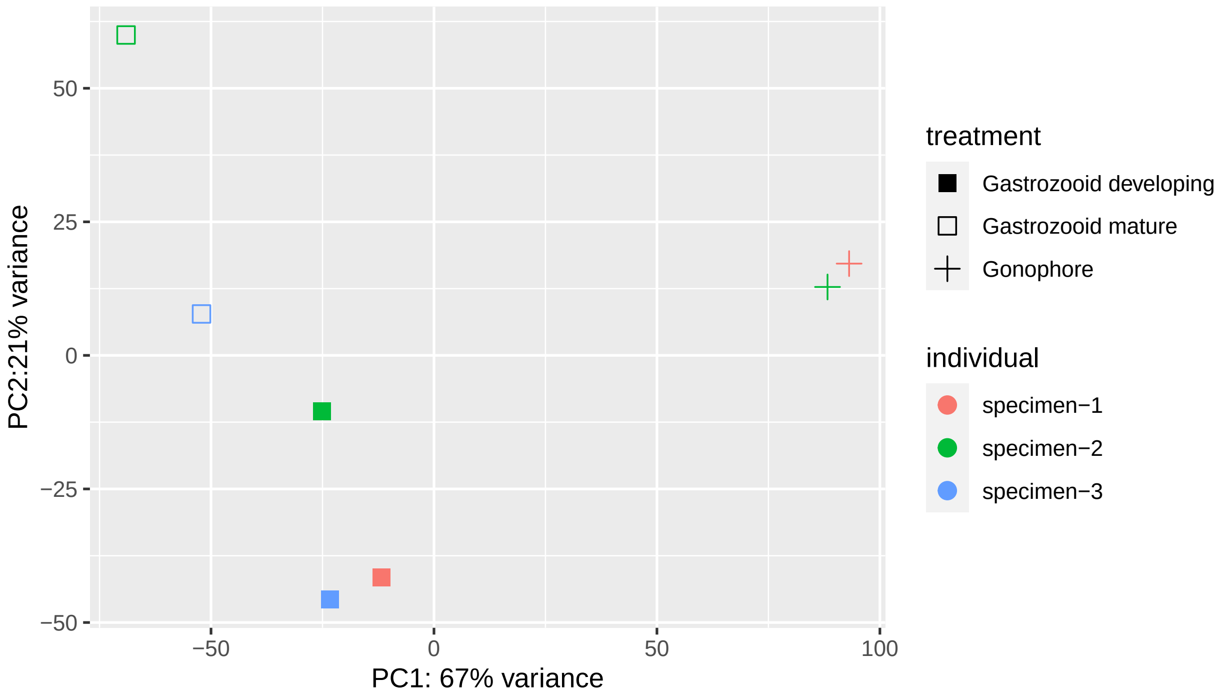


Figure S3: PCA of regularized log transformed expression counts of zooids/tissues in *Diphyes dispar*. Color indicates the replicate number, shape indicates the zooid.

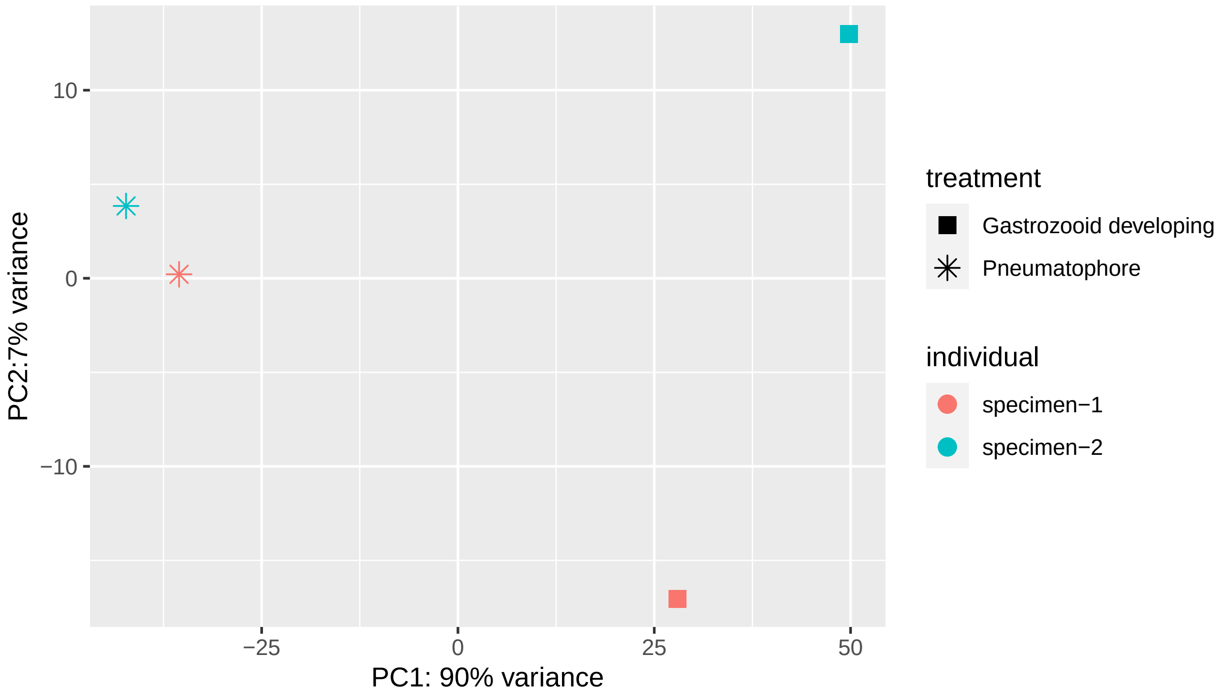


Figure S4: PCA of regularized log transformed expression counts of zooids/tissues in *Apolemia lanosa*. Color indicates the replicate number, shape indicates the zooid.

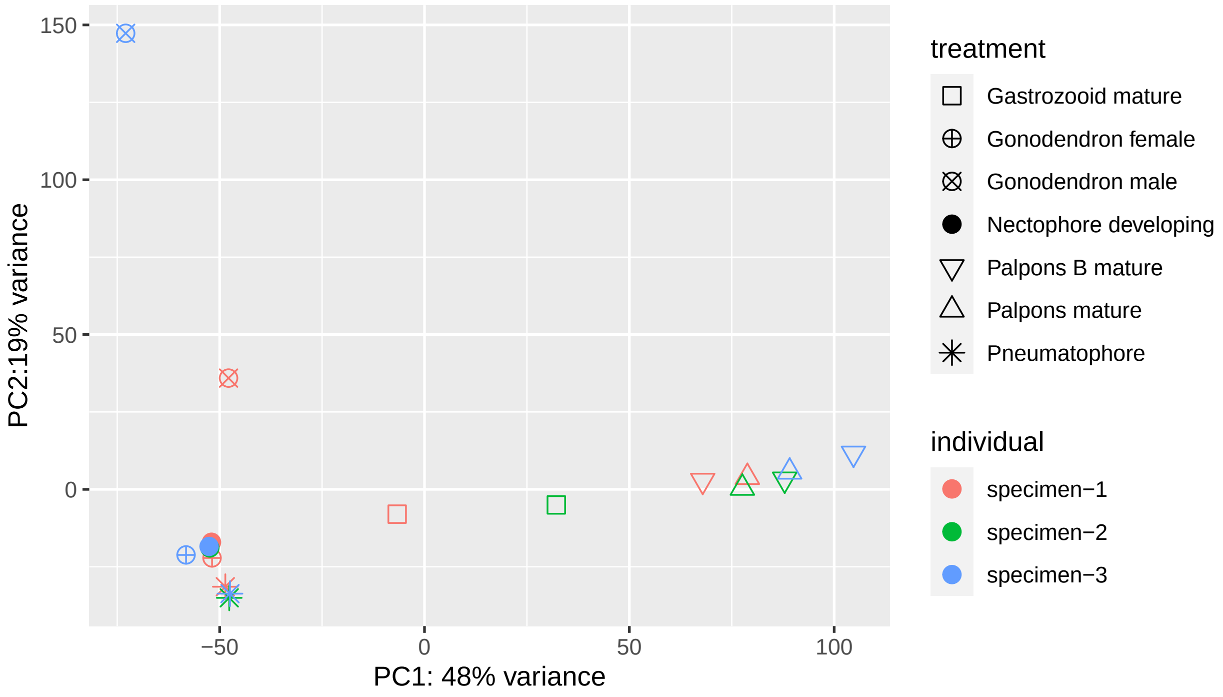


Figure S5: PCA of regularized log transformed expression counts of zooids/tissues in *Agalma elegans*. Color indicates the replicate number, shape indicates the zooid.

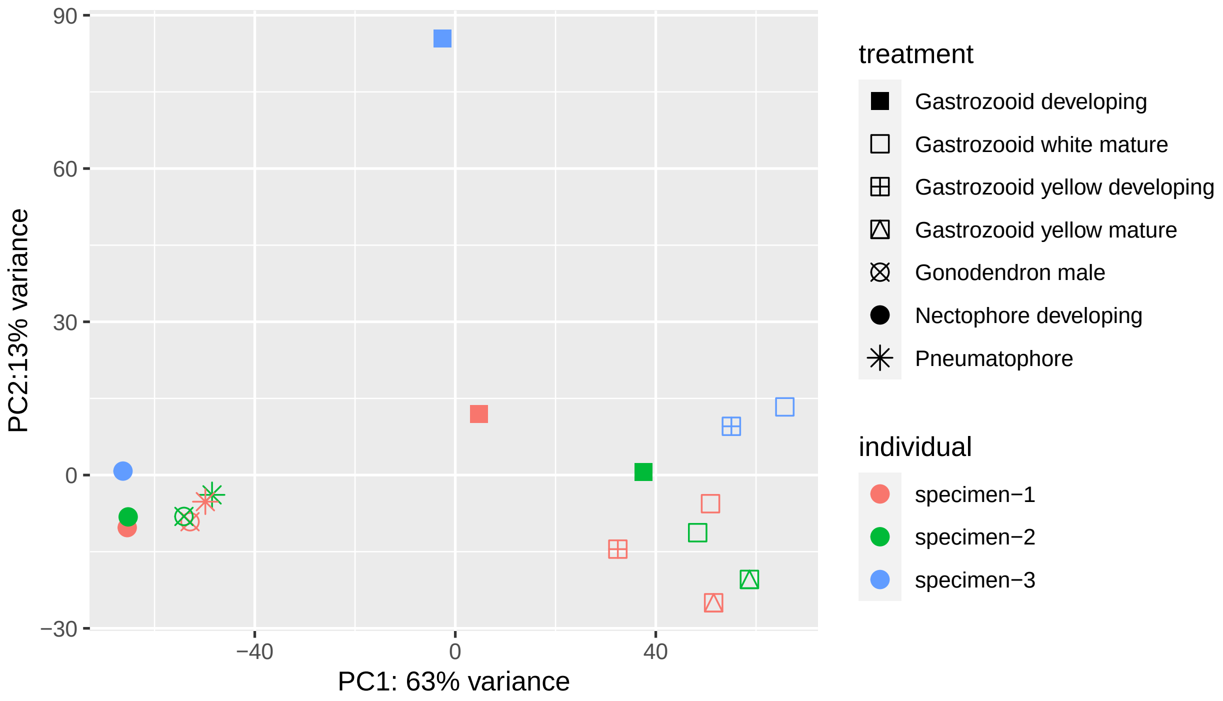


Figure S6: PCA of regularized log transformed expression counts of zooids/tissues in *Bargmannia elongata*. Color indicates the replicate number, shape indicates the zooid.

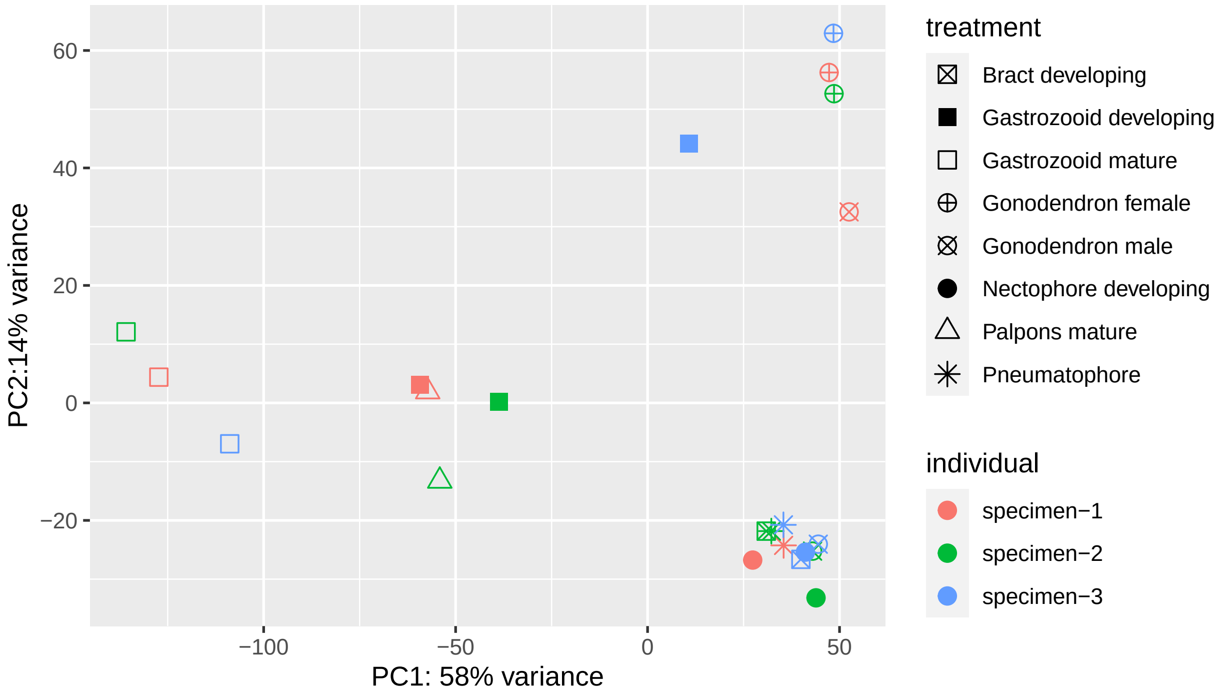


Figure S7: PCA of regularized log transformed expression counts of zooids/tissues in *Frillagalma vityazi*. Color indicates the replicate number, shape indicates the zooid.

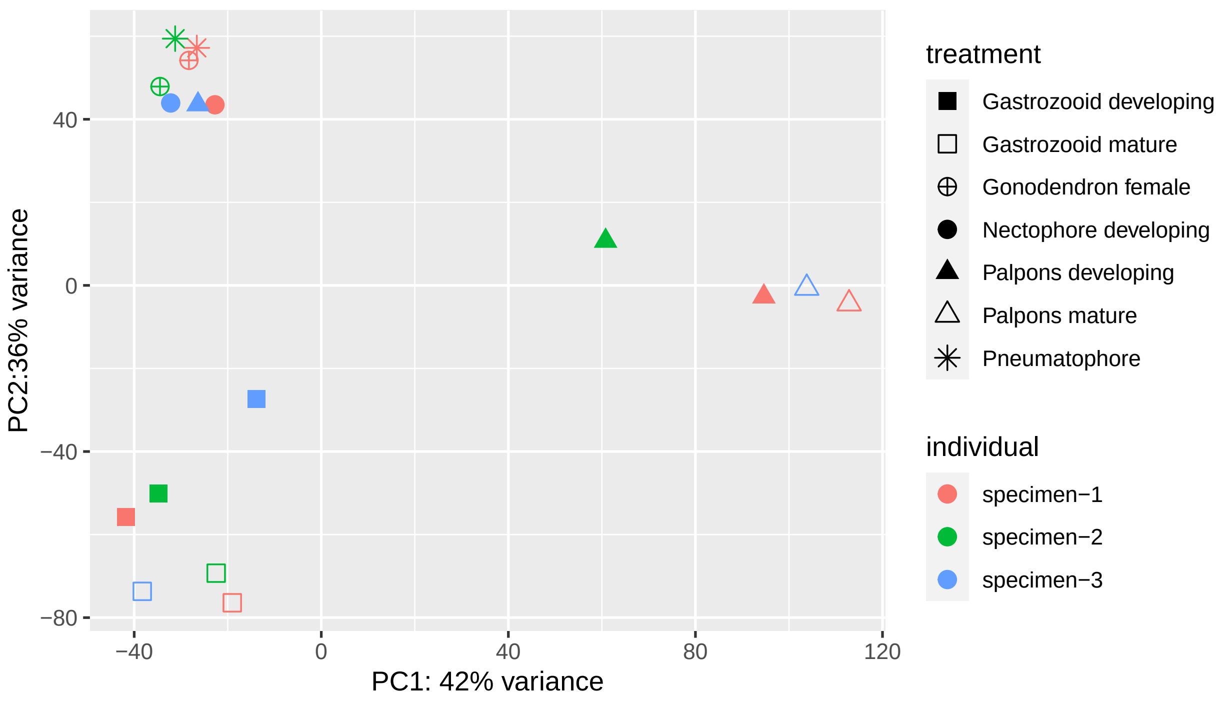


Figure S8: PCA of regularized log transformed expression counts of zooids/tissues in *Nanomia bijuga*. Color indicates the replicate number, shape indicates the zooid.

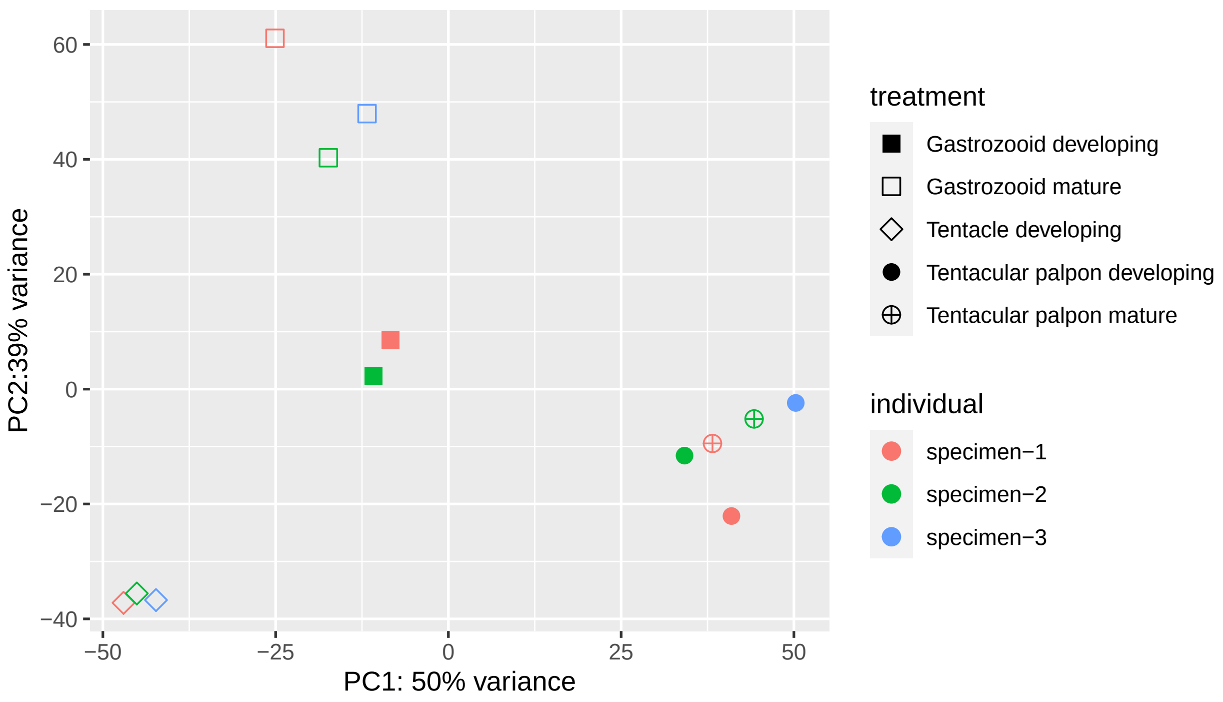


Figure S9: PCA of regularized log transformed expression counts of zooids/tissues in *Physalia physalis*. Color indicates the replicate number, shape indicates the zooid.

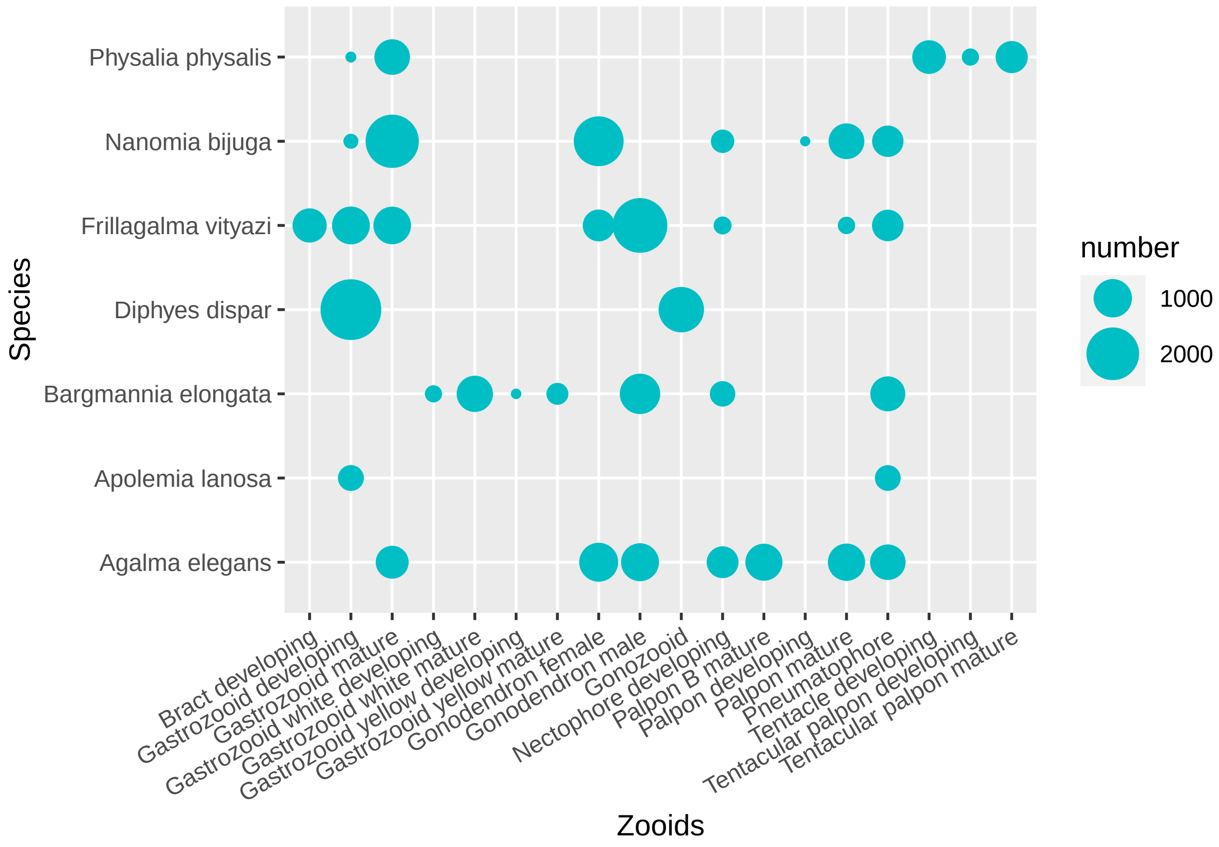


Figure S10: Number of genes identified as uniquely differentially expressed in zooids within different siphonophore species. Unique in this case means that the genes are significantly differentially expressed (higher) in these zooids and are not found to be significantly differentially expressed in any other zooid within the same species. Note that larger numbers of genes are identified in species where fewer zooids were sampled, this is likely due to sampling differences, as opposed to biological differences among species.

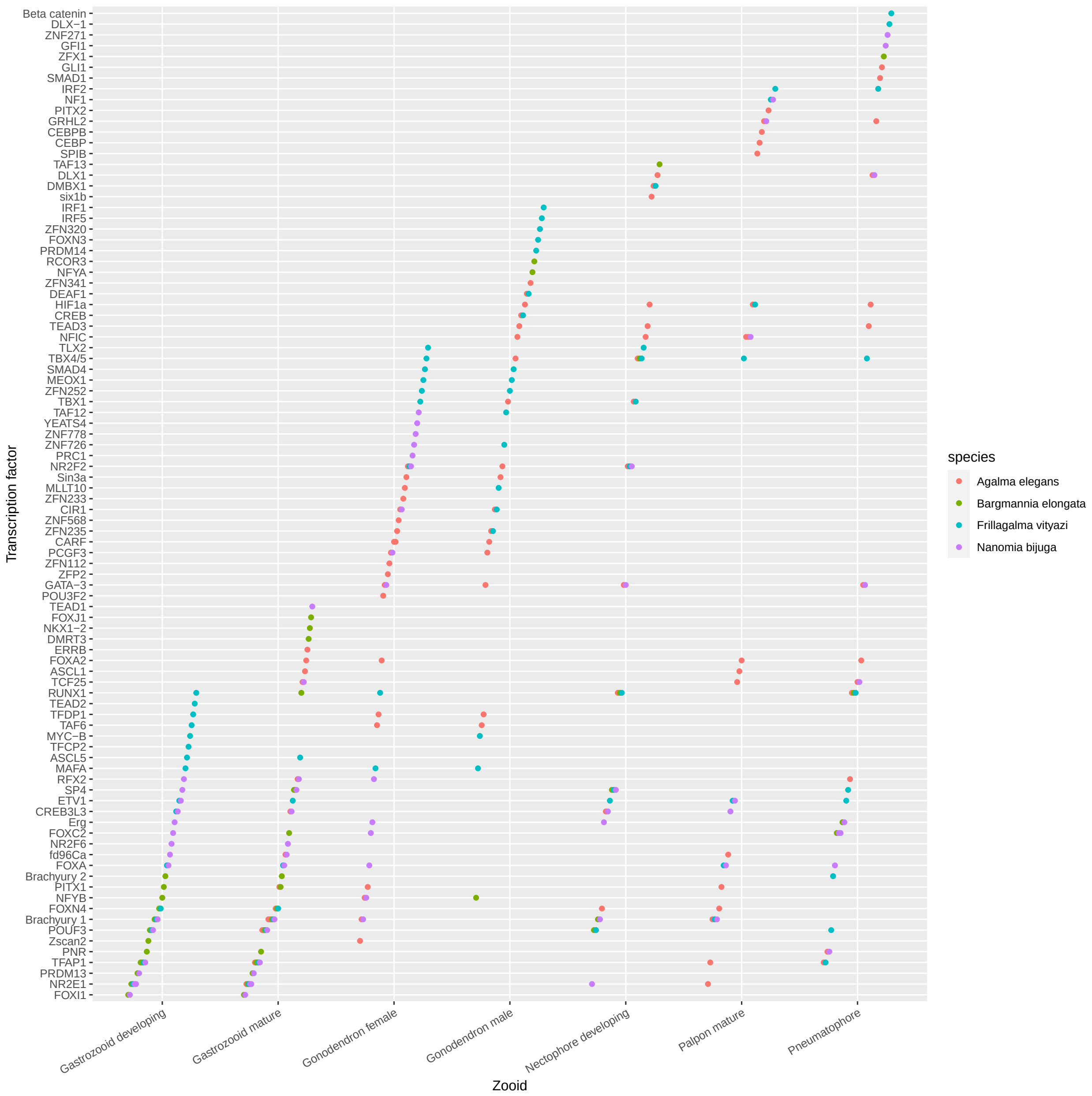


Figure S11: Putative transcription factors found to be significantly differentially expressed in zooids and the pneumatophore across four different siphonophore species. Transcription factor identity based on blast hit, GO DNA-binding transcription factor activity followed by manual curation. For *Bargmannia elongata* data is shown for 'white' gastrozooids.

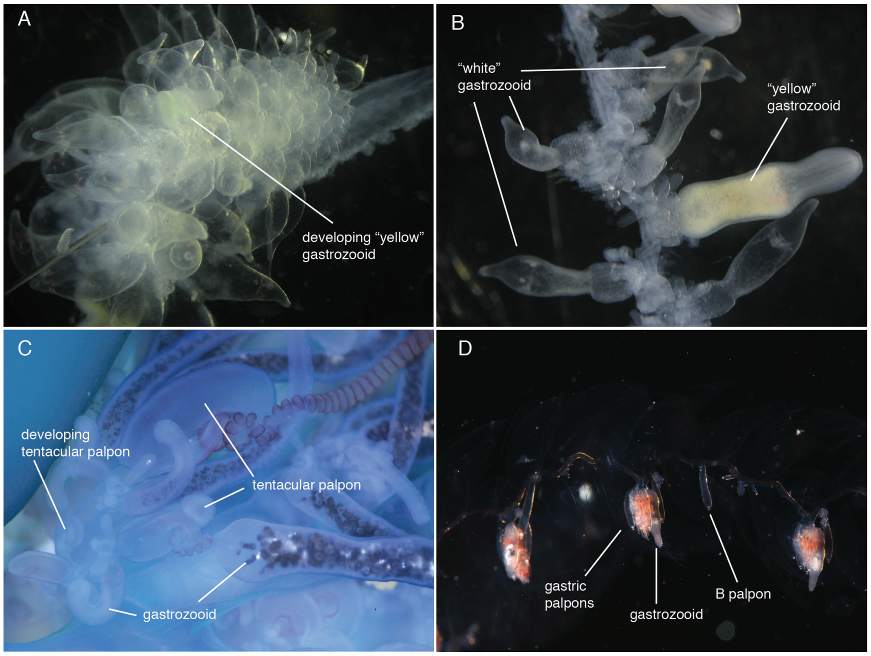


Figure S12: Unique siphonophore zooids that were sampled for differential gene expression. A. *Bargmannia elongata* with developing “yellow” gastrozooid surrounded by developing “white” gastrozooids. B. *Bargmannia elongata* stem with mature “white” and “yellow” gastrozooids. C. Zooids in *Physalia physalis*, including multiple developing stages of gastrozooids, tentacular palpon and tentacle; the most mature form of either zooid is not shown. D. Stem of *Agalma elegans*, with gastric palpons, B-palpon and gastrozooid shown.

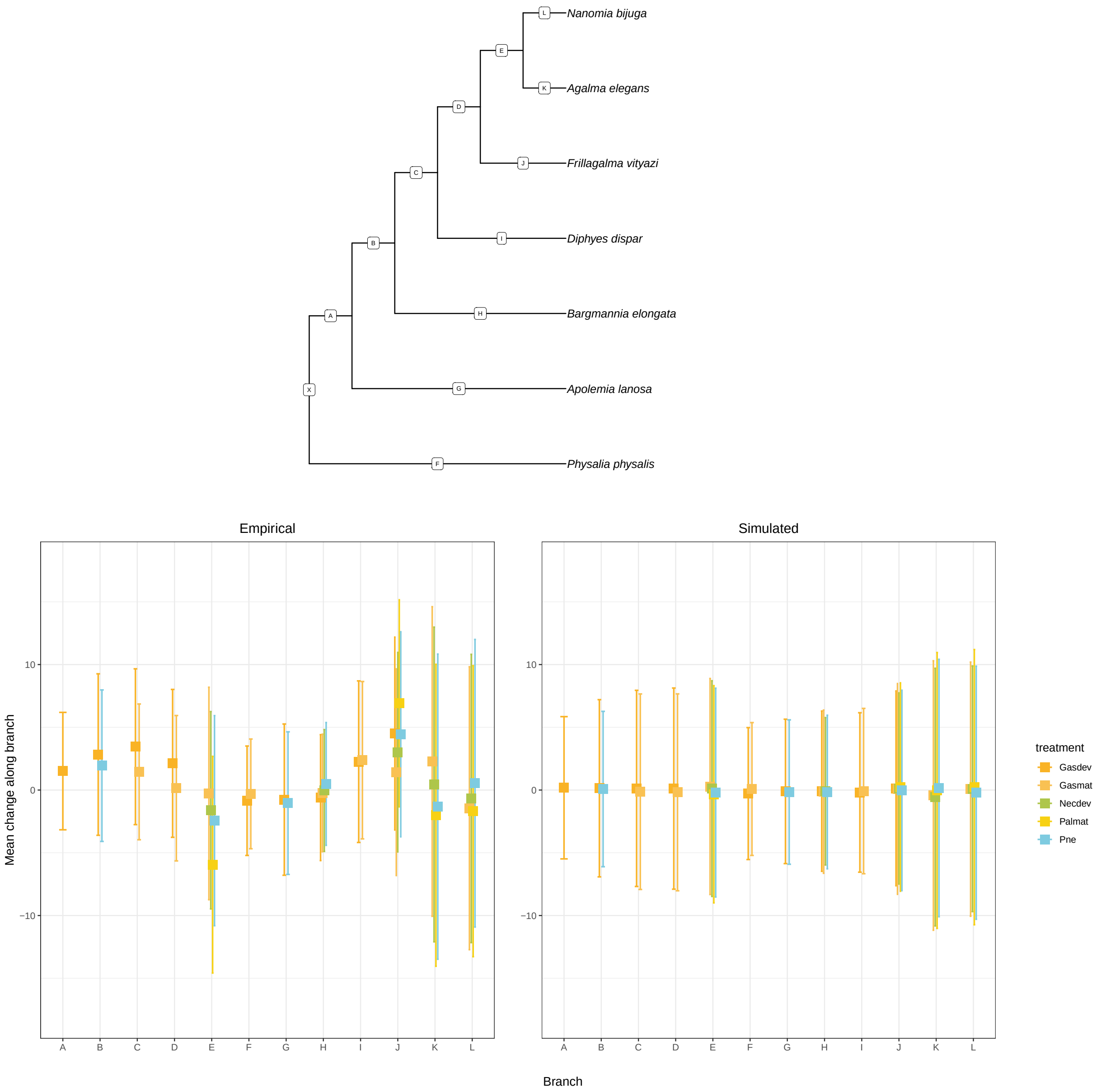


Figure S13: Mean changes of TPM10K along branches across all gene trees that correspond to branches in the species trees, error bar is one standard deviation. Top panel: species phylogeny with branch IDs given as letters. Lower panel: Distribution of changes along a species-equivalent branch in a gene tree, showing mean change in the empirical dataset and in the BM simulation. Treatment type is coded in colour, Gasdev = Developing Gastrozooid, Gasmat= Mature gastrozooid, Necdev= Developing nectophore, Palmat = Mature palpon, Pne= pneumatophore.

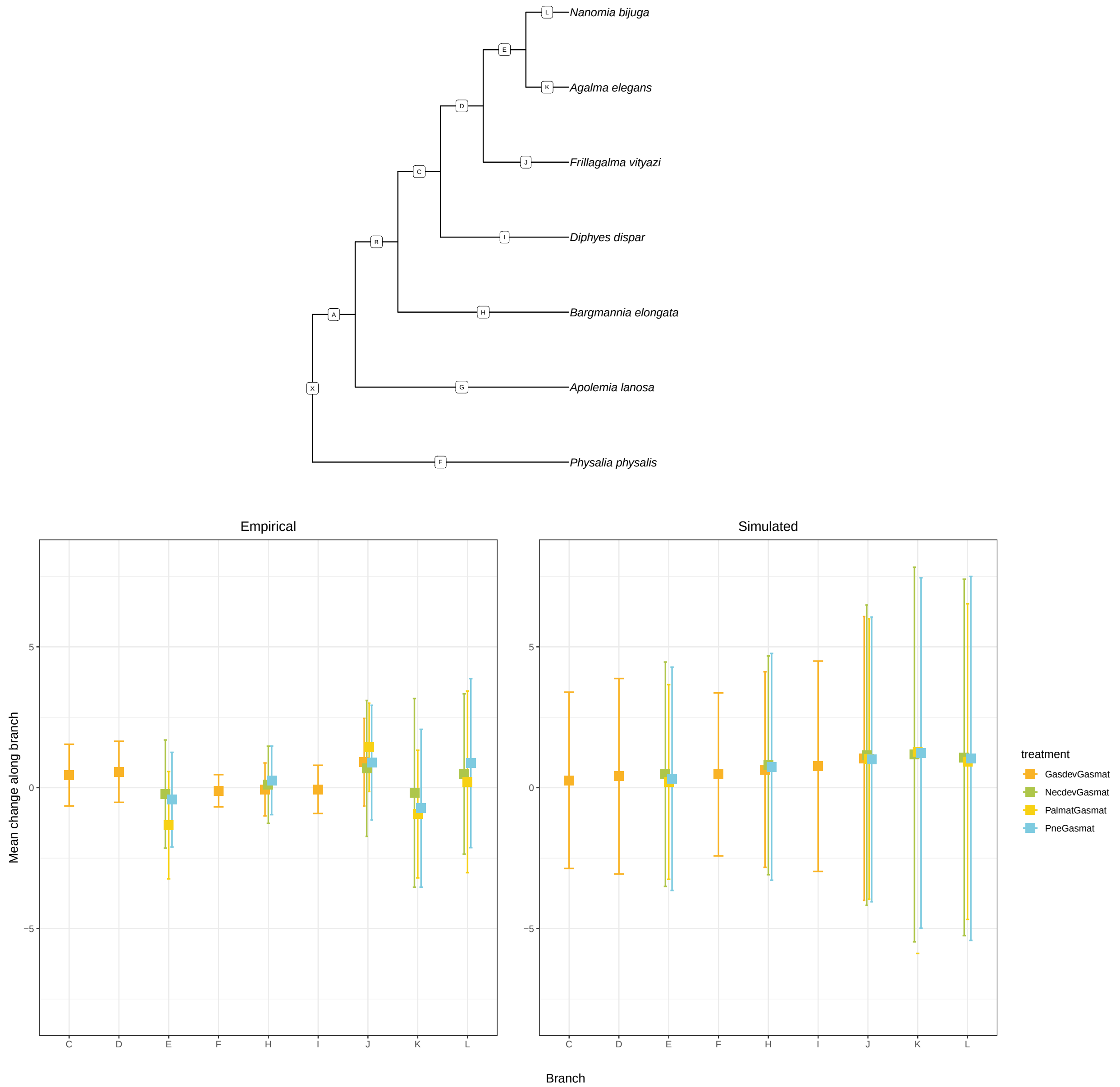


Figure S14: Mean changes of expression ratios along branches across all gene trees that correspond to branches in the species tree, error bar is one standard deviation. Top panel: species phylogeny with branch IDs given as letters. Lower panel: Distribution of changes along a branch in a gene tree, showing mean change in the empirical dataset and in the BM simulation. Treatment type is coded in colour, GasdevGasmat = developing gastrozooid / mature gastrozooid, NecdevGasmat = developing nectophore / mature gastrozooid, PalmatGasmat = mature palpon / mature gastrozooid, PneGasmat = Pneumatophore / mature gastrozooid.

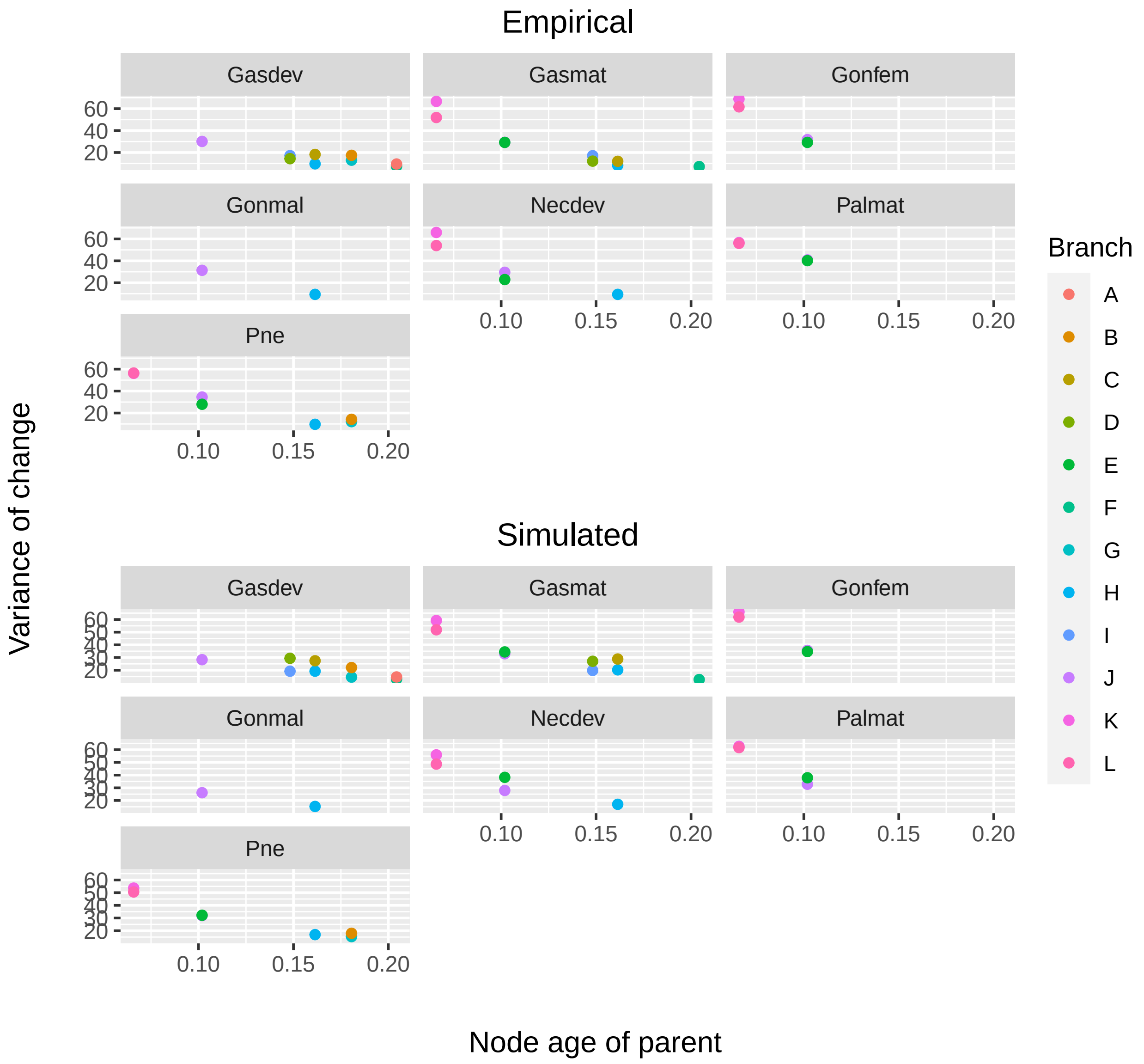


Figure S15: Variance of TPM10K change across a branch plotted against the age of the parent node. Branch ID is coded in colour, values are separated by treatment. Gasmat = Mature Gastrozooid, Gasdev = Developing Gastrozooid, Necdev= Developing nectophore, Palmat = Mature palpon , Pne= Pneumatophore

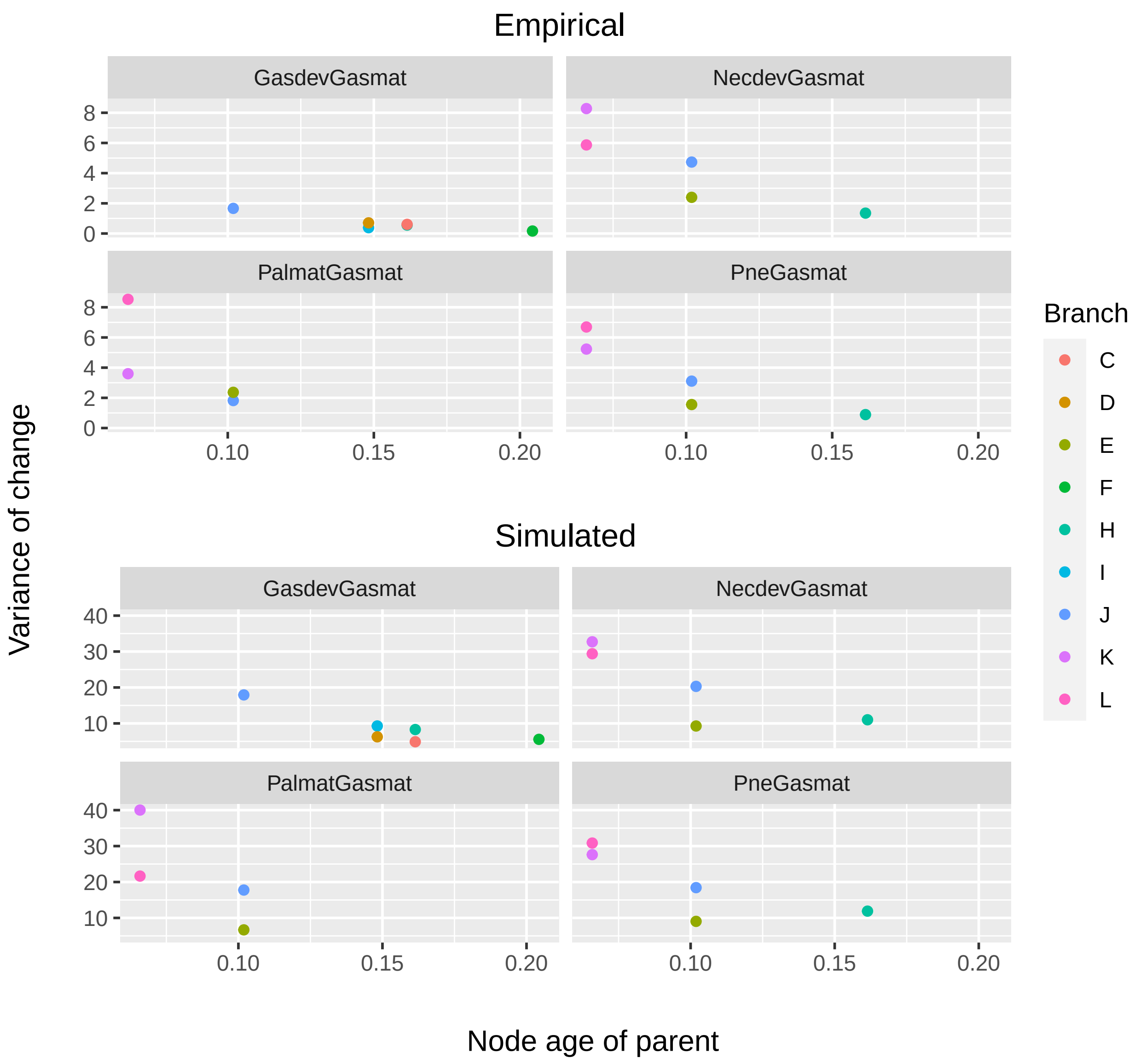


Figure S16: Variance of expression ratio change across a branch plotted against the age of the parent node. Branch ID is coded in colour, values are separated by treatment. GasdevGasmat = Developing Gastrozooid vs Mature Gastrozooid, NecdevGasmat= Developing nectophore vs Mature Gastrozooid, PalmatGasmat = Mature palpon vs Mature Gastrozooid, PneGasmat= Pneumatophore vs Mature Gastrozooid

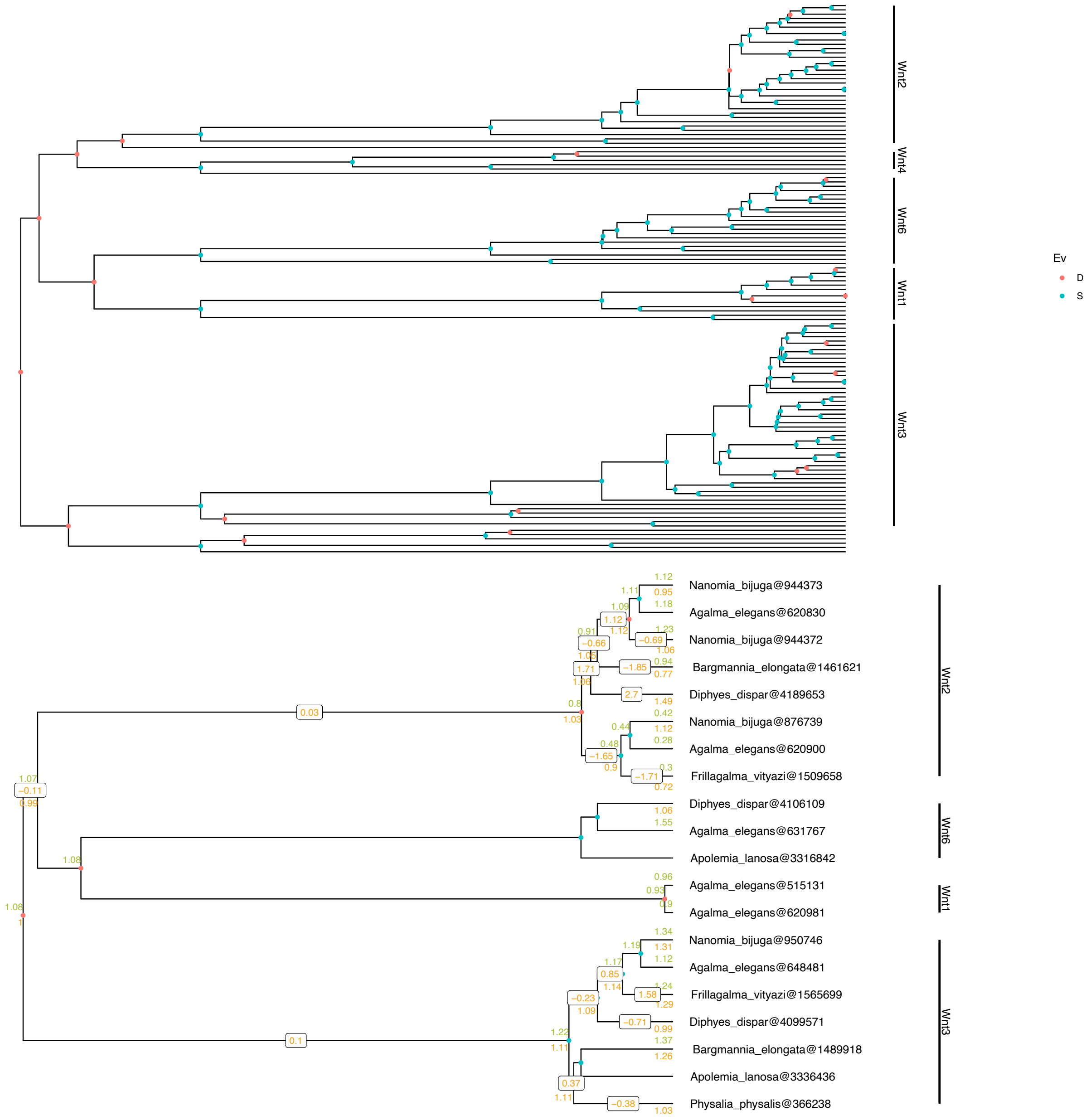


Figure S17: Top panel: Maximum likelihood gene tree of the Wnt family in Cnidaria, Wnt clades that include siphonophore sequences are highlighted. Bottom panel: Pruned gene tree of wnt genes in siphonophore species with expression values (TPM10K) at the tips and nodes. Developing nectophore/mature gastrozooid expression in green (above node), developing gastrozooid/mature gastrozooid expression in orange (below node). Changes across the branch are shown for developing gastrozooid/mature gastrozooid (orange). Red circles are duplication nodes and blue circles are speciation nodes.

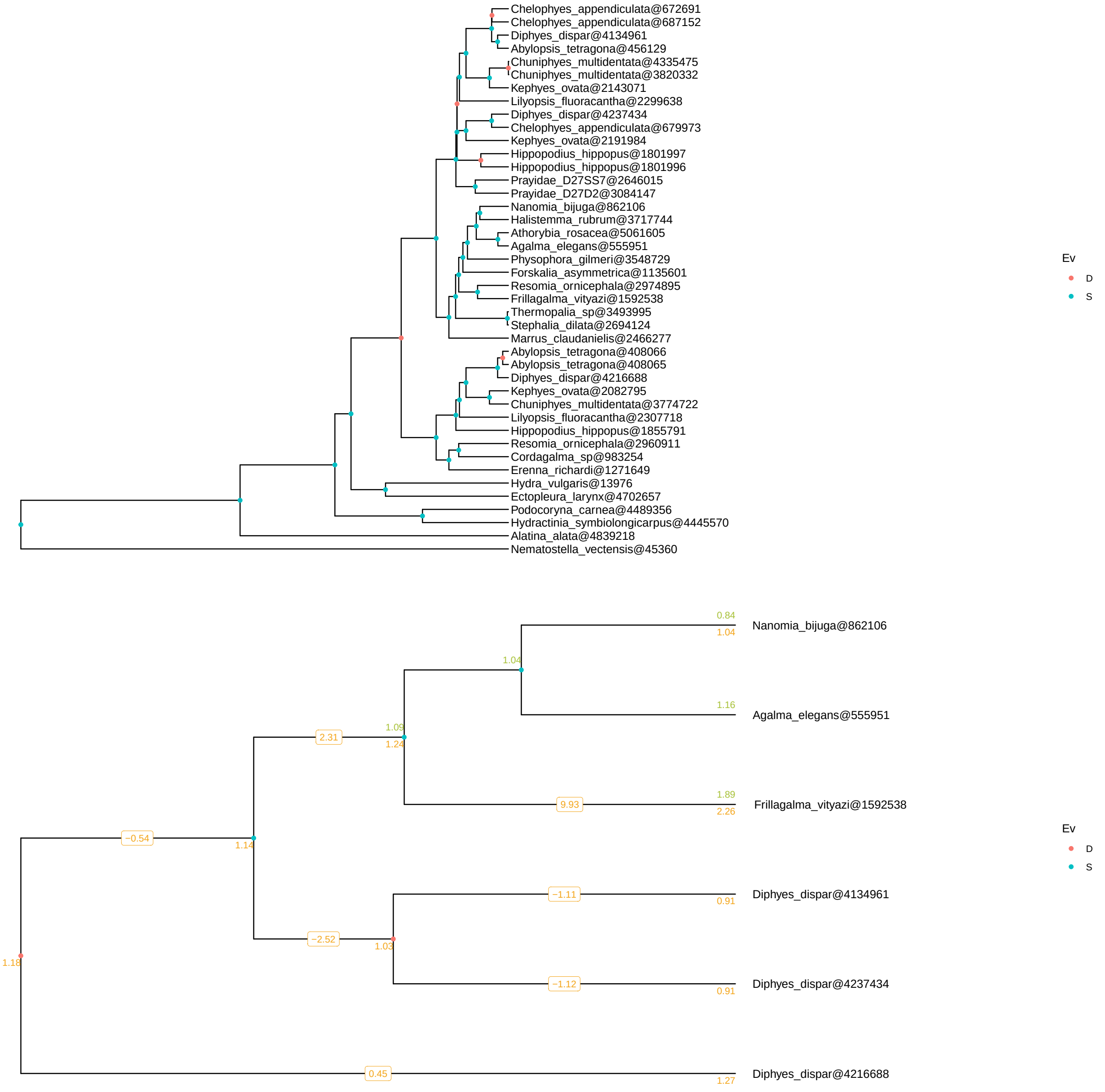


Figure S18: Top panel: Maximum likelihood gene tree of the Wnt4 in Cnidaria. Bottom panel: Pruned gene tree of wnt4 genes in siphonophore species with expression values (TPM10K) at the tips and nodes. Developing nectophore/mature gastrozooid expression in green (above node), developing gastrozooid/mature gastrozooid expression in orange (below node). Changes across the branch are shown for developing gastrozooid/mature gastrozooid (orange). Red circles are duplication nodes and blue circles are speciation nodes.

**Software versions**

This manuscript was computed on Thu Jul 29 18:23:02 2021 with the following R package versions.

```
R version 4.0.0 (2020-04-24)
Platform: x86_64-apple-darwin17.0 (64-bit)
Running under: macOS Catalina 10.15.7

Matrix products: default
BLAS:   /Library/Frameworks/R.framework/Versions/4.0/Resources/lib/libRblas.dylib
LAPACK: /Library/Frameworks/R.framework/Versions/4.0/Resources/lib/libRlapack.dylib

locale:
[1] en_US.UTF-8/en_US.UTF-8/en_US.UTF-8/C/en_US.UTF-8/en_US.UTF-8

attached base packages:
 [1] grid      parallel  stats4    stats     graphics  grDevices utils
 [8] datasets  methods   base

other attached packages:
 [1] gtools_3.8.2                ggrepel_0.8.2
 [3] kableExtra_1.1.0            corrplot_0.84
 [5] ggpubr_0.3.0                lme4_1.1-23
 [7] FSA_0.8.30                  emmeans_1.4.6
 [9] gridExtra_2.3               geiger_2.0.6.4
[11] GGally_1.5.0                PhylogeneticEM_1.4.0
[13] Matrix_1.2-18               magrittr_1.5
[15] digest_0.6.25               forcats_0.5.0
[17] stringr_1.4.0               dplyr_0.8.5
[19] purrr_0.3.4                 readr_1.3.1
[21] tidyr_1.0.3                 tibble_3.0.1
[23] tidyverse_1.3.0             RColorBrewer_1.1-2
[25] picante_1.8.1               nlme_3.1-147
[27] vegan_2.5-6                 lattice_0.20-41
[29] permute_0.9-5               ape_5.3
[31] fields_10.3                 maps_3.3.0
[33] spam_2.5-1                  dotCall64_1.0-0
[35] knitr_1.28                  bookdown_0.19
[37] ggplot2_3.3.0               agalmar_0.0.0.9000
[39] hutan_0.5.1                 vsn_3.56.0
[41] goseq_1.40.0                geneLenDataBase_1.24.0
[43] BiasedUrn_1.07              topGO_2.40.0
[45] SparseM_1.78                GO.db_3.11.1
[47] AnnotationDbi_1.50.0        graph_1.66.0
[49] DESeq2_1.28.1               SummarizedExperiment_1.18.1
[51] DelayedArray_0.14.0         matrixStats_0.56.0
[53] Biobase_2.48.0              GenomicRanges_1.40.0
[55] GenomeInfoDb_1.24.0         IRanges_2.22.1
[57] S4Vectors_0.26.0            BiocGenerics_0.34.0
[59] impute_1.62.0               treeio_1.12.0
[61] ggtree_2.2.1

loaded via a namespace (and not attached):
  [1] tidyselect_1.1.0        RSQLite_2.2.0           BiocParallel_1.22.0
  [4] munsell_0.5.0           codetools_0.2-16        preprocessCore_1.50.0
```

```
  [7] statmod_1.4.34          withr_2.2.0             colorspace_1.4-1
 [10] rstudioapi_0.11         ggsignif_0.6.0          labeling_0.3
 [13] GenomeInfoDbData_1.2.3  farver_2.0.3            bit64_0.9-7
 [16] coda_0.19-3             vctrs_0.3.0             generics_0.0.2
 [19] xfun_0.13               BiocFileCache_1.12.0    R6_2.4.1
 [22] locfit_1.5-9.4          bitops_1.0-6            reshape_0.8.8
 [25] assertthat_0.2.1        scales_1.1.1            gtable_0.3.0
 [28] affy_1.66.0             rlang_0.4.6             genefilter_1.70.0
 [31] splines_4.0.0           rtracklayer_1.48.0      rstatix_0.5.0
 [34] lazyeval_0.2.2          broom_0.5.6             reshape2_1.4.4
 [37] abind_1.4-5             BiocManager_1.30.10     yaml_2.2.1
 [40] modelr_0.1.7            GenomicFeatures_1.40.0  backports_1.1.7
 [43] tools_4.0.0             affyio_1.58.0           ellipsis_0.3.0
 [46] Rcpp_1.0.4.6            plyr_1.8.6              progress_1.2.2
 [49] zlibbioc_1.34.0         RCurl_1.98-1.2          prettyunits_1.1.1
 [52] openssl_1.4.1           deSolve_1.28            haven_2.2.0
 [55] cluster_2.1.0           fs_1.4.1                data.table_1.12.8
 [58] openxlsx_4.1.5          reprex_0.3.0            mvtnorm_1.1-0
 [61] hms_0.5.3               patchwork_1.0.0         evaluate_0.14
 [64] xtable_1.8-4            XML_3.99-0.3            rio_0.5.16
 [67] readxl_1.3.1            compiler_4.0.0          biomaRt_2.44.0
 [70] crayon_1.3.4            minqa_1.2.4             htmltools_0.4.0
 [73] mgcv_1.8-31             geneplotter_1.66.0      aplot_0.0.4
 [76] lubridate_1.7.8         DBI_1.1.0               dbplyr_1.4.3
 [79] subplex_1.6             MASS_7.3-51.6           rappdirs_0.3.1
 [82] boot_1.3-25             car_3.0-7               cli_2.0.2
 [85] pkgconfig_2.0.3         rvcheck_0.1.8           GenomicAlignments_1.24.0
 [88] foreign_0.8-79          xml2_1.3.2              foreach_1.5.0
 [91] annotate_1.66.0         webshot_0.5.2           XVector_0.28.0
 [94] estimability_1.3        rvest_0.3.5             Biostrings_2.56.0
 [97] rmarkdown_2.1           cellranger_1.1.0        tidytree_0.3.3
[100] curl_4.3                Rsamtools_2.4.0         nloptr_1.2.2.1
[103] lifecycle_0.2.0         jsonlite_1.6.1          carData_3.0-3
[106] viridisLite_0.3.0       askpass_1.1             limma_3.44.1
[109] fansi_0.4.1             pillar_1.4.4            httr_1.4.1
[112] survival_3.1-12         glue_1.4.1              zip_2.0.4
[115] iterators_1.0.12        bit_1.1-15.2            stringi_1.4.6
[118] blob_1.2.1              memoise_1.1.0
```
